# FTO does not function as an m^6^A eraser in AML: a reassessment using direct RNA nanopore sequencing

**DOI:** 10.1101/2025.10.22.681652

**Authors:** Luke S. Nicholson, Catarina Guimarães-Teixeira, Jianheng Fox Liu, Raymond Chien, Jin Xu, Hui Xian Poh, Ying-Xian Pan, Samie R. Jaffrey

## Abstract

A central concept in epitranscriptomics is that the RNA demethylase FTO acts as a site-selective eraser of *N*^6^-methyladenosine (m^6^A) in specific mRNAs. This is most well-established in acute myeloid leukemia (AML), where demethylation of *MYC* is thought to promote leukemogenesis. Using nanopore direct RNA sequencing to quantify m^6^A transcriptome-wide, we find that m^6^A stoichiometry, including at *MYC*-specific sites, remains unaffected by *FTO* knockdown or pharmacologic inhibition in AML. Conversely, levels of *N*^6^-2’-O-dimethyladenosine (m^6^Am) are markedly increased in snRNAs, indicating that m^6^Am, not m^6^A, is the major substrate of FTO in AML. We extend these results outside the AML context in *FTO* knockout brain and HEK293T cells, detecting no selective increases in mRNA m^6^A but widespread increases in snRNA m^6^Am levels following FTO depletion. These findings suggest that FTO has minimal ability to erase m^6^A in mRNA. Instead, the reported biological effects of FTO are likely mediated by demethylation of m^6^Am.

## INTRODUCTION

*N*^6^-methyladenosine (m^6^A) is the most abundant internal modification of eukaryotic mRNA. m^6^A plays a critical role in regulating mRNA stability, and can affect translation, splicing, and other aspects of RNA processing^1^. A major concept in the epitranscriptomics field is that m^6^A is a dynamic and reversible modification. This idea was introduced as a result of the discovery that the fat mass and obesity-associated protein (FTO) acts as an m^6^A demethylase^2^. The initial studies on FTO used *in vitro* assays to demonstrate that FTO can remove the methyl group of m^6^A in RNA oligonucleotides, converting m^6^A to A. This *in vitro* finding was supported by mass spectrometry analysis of FTO-depleted HeLa and HEK293FT cells, which showed a small but significant increase in m^6^A levels in cellular mRNA.

Subsequently, seminal studies in acute myeloid leukemia (AML) provided the critical concept that FTO is a transcript-specific demethylase. AML cells were proposed to be particularly relevant for FTO-mediated demethylation of m^6^A in mRNA because FTO was reported to be cytosolic in these cells, in contrast to its nuclear localization in most other cell types^3^. In this way, FTO would be much more effective at demethylating m^6^A on mRNA in AML cell lines. The initial study by Li et al. reported that FTO functions as an oncogene in AML and promotes leukemogenesis by demethylating m^6^A on key transcripts, including *ASB2* and *RARA*^4^. Building on this, Su et al. showed that FTO demethylates m^6^A sites in *MYC*, a master transcriptional regulator in cancer. Demethylation of m^6^A sites in *MYC* leads to stabilization of *MYC* mRNA and increased MYC protein expression. They also showed that FTO activity can be regulated by the oncometabolite R-2-hydroxyglutarate, which inhibits FTO, leading to increased m^6^A levels in *MYC* and impaired AML cell growth^5^.

Because of the importance of FTO in regulating key cancer-related transcripts, Huang et al. developed FB23-2, a small-molecule FTO inhibitor that increased m^6^A levels on *MYC* and other AML-relevant transcripts and demonstrated potent anti-leukemic effects in both cell lines and xenograft models^6^. Thus, FTO has a key oncogenic role in AML by controlling the m^6^A levels in critical cancer-driving transcripts^5^.

These seminal studies led to a wave of subsequent research that identified additional mRNA targets of FTO in different cancers and other disease contexts^7–9^. Overall, these results have led to the widely accepted model that specific m^6^A sites in specific mRNAs are erased by FTO, providing a basis for reversible and dynamic regulation of m^6^A levels.

Other studies have raised questions about the biological relevance of FTO-mediated m^6^A demethylation^10^. Although m^6^A can clearly be demethylated by FTO *in vitro*, we noted that the catalytic rate was exceptionally slow compared to the catalytic rates of similar demethylases acting on their cognate substrates^10^. Additionally, FTO-mediated m^6^A demethylation required a large amount of FTO enzyme relative to the m^6^A substrate^2^. We tested other possible substrates and found that another modified nucleotide, *N*^6^, 2’-O-dimethyladenosine (m^6^Am), was demethylated with a ∼100-fold higher catalytic efficiency than m^6^A and was efficiently demethylated with low concentrations of FTO^10^. m^6^Am is located at the first transcribed nucleotide of snRNAs, snoRNAs, and mRNAs^10–12^ and is structurally similar to m^6^A but contains a methyl modification on the 2’-hydroxyl of the ribose. Using MeRIP-seq, a transcriptome-wide m^6^A mapping method developed by us and others^13,14^, we found no clear increases in m^6^A sites in any mRNA in *FTO* knockout brain^15^. However, we observed clear increases in *N*^6^,2′-*O*-dimethyladenosine (m^6^Am) in snRNA in FTO-depleted HEK293T cells^11,12^, with smaller effects on m^6^Am in mRNA^10,11^.

Despite these findings, the widespread view is that the ability of FTO to demethylate m^6^A is dependent on cell type, with AML cell lines being particularly well-suited to FTO-mediated m^6^A demethylation due to the high levels of cytosolic FTO^3^. This conclusion is supported by a large body of literature that used m^6^A mapping methods to compare the transcriptome-wide landscape of m^6^A in control and FTO-deficient cells. Each of these studies reported specific transcripts that are efficiently demethylated by FTO. An important concern is that these studies relied almost exclusively on MeRIP-seq. MeRIP-seq uses m^6^A antibodies to immunoprecipitate m^6^A-containing RNA fragments for sequencing^13,14^. Unless spike-in standards or simultaneous pooling methods (e.g., m6A-seq2) are used, the resulting peaks can exhibit considerable variability in peak heights, even between biological replicates^16,17^. This occurs because antibody binding and RNA precipitation efficiency are highly sensitive to subtle differences in washing and other immunoprecipitation conditions^16^. Since the specific mRNA targets of FTO were identified based on differential peak heights that occurred after FTO depletion, these peak height differences might not reflect authentic changes in m^6^A levels. Instead, these putative FTO-regulated m^6^A sites could simply be the normal peak height variability in MeRIP-seq.

In order to definitively establish that an m^6^A site is physiologically demethylated by FTO, the exact stoichiometry of the m^6^A site needs to be established in the presence and absence of FTO. An m^6^A site that is normally demethylated by FTO should have low stoichiometry under control conditions and increased m^6^A stoichiometry once FTO is depleted. Since MeRIP-seq does not provide single-nucleotide resolution and is not quantitative, the magnitude of the stoichiometry changes in methylation achieved by FTO depletion could not be reported in the AML studies or in subsequent studies of FTO and m^6^A.

To identify the mRNAs that contain m^6^A sites that are regulated by FTO, we used Oxford Nanopore direct RNA sequencing, which provides highly accurate measurements of m^6^A stoichiometry at individual sites throughout the transcriptome^18^. We measured m^6^A sites in the AML cell lines MONOMAC-6 and MOLM-13 under conditions of *FTO* knockdown and pharmacologic FTO inhibition. Contrary to expectations based on the published studies, we did not observe any significant changes in m^6^A stoichiometry in *MYC* or any other proposed FTO targets, or any m^6^A sites at all following FTO depletion in AML cell lines. We also assessed other FTO-depleted tissues. In Fto-depleted mouse brain samples, we found no evidence for increased m^6^A stoichiometry at any sites. In FTO-depleted HEK293T cells, we found subtle increases in global m^6^A stoichiometry but no evidence for selectivity in demethylation by FTO. These results indicate that FTO does not target specific mRNAs and, at best, has minimal effect on m^6^A in cells. Overall, our data highlight the importance of re-examining FTO’s proposed ‘m^6^A eraser’ role using quantitative m^6^A mapping methods.

## RESULTS

### Nanopore direct RNA sequencing provides highly accurate m^6^A stoichiometry measurements

Recent improvements in the nanopore technology and flow cell chemistry (“RNA004”) have allowed m^6^A to be detected in single RNA reads with high accuracy^19,20^. m^6^A is detected using the Dorado basecaller, a neural network trained on raw nanopore current signals to assign per-read modification probabilities^21^. When multiple independent mRNAs for a specific gene are sequenced, the stoichiometry of m^6^A at any site within the mRNA can be calculated^21–24^.

The accuracy of m^6^A stoichiometry measurements by nanopore sequencing has been established by comparing stoichiometry measurements obtained using GLORI^25^, a highly accurate and quantitative m^6^A mapping method that uses a chemical approach to distinguish A from m^6^A on a transcriptome-wide level^26–28^. In our hands, we similarly found a very high correlation between m^6^A stoichiometry predicted by Dorado and stoichiometries measured by GLORI in HEK293T cells^25^ (**Figure S1A**). A benefit of nanopore sequencing is that it also detects m^6^A in long reads, which is not possible with GLORI.

To further examine the reliability of nanopore direct RNA sequencing, we examined m^6^A stoichiometries in cells lacking METTL3, the primary methyltransferase responsible for m^6^A formation in mRNA^29,30^. We performed nanopore direct RNA sequencing on RNA from wild-type and *Mettl3* knockout mouse embryonic stem cells (mESCs)^30^ and plotted the site-specific stoichiometries. As expected, all sites in the knockout exhibited stoichiometries of zero or near zero (**Figure S1B**). These controls indicate that, in our hands and under the stated parameters, nanopore direct RNA sequencing provides accurate m^6^A stoichiometry estimates with a low false-positive rate.

### Establishing the variability of m^6^A stoichiometry measurements between nanopore replicates

Before analyzing FTO-depleted cells, we established the baseline sample-to-sample variability of nanopore m^6^A measurements. Because m^6^A stoichiometry calculations are inherently subject to statistical sampling noise, small stoichiometric differences between identical biological replicates will inevitably occur. To determine the sequencing depth required to sufficiently suppress this noise, we tested different minimum read coverage thresholds using two independent control conditions (DMSO-treated and non-targeting shRNA-treated MONOMAC-6 cells). As expected, evaluating control-to-control concordance revealed that higher read depths rapidly increased the Pearson’s correlation (R²) and progressively tightened both the limits of agreement and the cumulative distribution of per-site stoichiometry differences between samples (Δm^6^A) (**Figure S1C–S1E**). We selected a 50-read minimum for all downstream analyses, as it provided the optimal balance between minimizing sampling variance and retaining a broad, transcriptome-wide dataset (**Figure S1F**).

Applying this 50-read threshold, we compared m^6^A stoichiometries across the control conditions to establish a baseline for identifying FTO-regulated sites. While the datasets correlated highly overall (Pearson’s R² = 0.959; **Figure S1G**), our primary focus was on the stochastic scatter *around* this correlation line, representing meaningless technical variance. A Bland-Altman analysis established that at a 50-read depth, 95% of site-level stoichiometry differences between independent controls fall within a band of ±0.124 (**Figure S1H**). Applying this thresholding to the MOLM-13 control datasets, we similarly see a range of ±0.123 (**Figure S1I**). By defining ±0.124 as the expected limit of statistical sampling noise at this specific read depth, we established a quantitative threshold: any apparent change in m^6^A stoichiometry following FTO depletion must exceed this margin to be considered a biologically meaningful difference.

### FTO depletion leads to marked upregulation of m^6^Am in AML snRNA

To directly measure the effect of FTO depletion on m^6^A and m^6^Am stoichiometries, we treated MONOMAC-6 and MOLM-13 cells with an FTO-specific shRNA or non-targeting shRNA lentivirus before proceeding with nanopore direct RNA sequencing, CROWN-seq, or GLORI (**Figure 1A**). Knockdown was used to deplete FTO since this is the primary method used in the studies of FTO regulation of m^6^A in AML cell lines^3–6^. Two lentiviruses were first tested in MONOMAC-6 cells (**Figure S1J**). shFTO-1 exhibited near-complete depletion of FTO protein and was thus used to test the role of FTO on m^6^A levels. This shRNA was the same one used in the prior studies of FTO in AML cell lines^4–6^. MONOMAC-6 was chosen as our primary cell line of study, as it was also used in the initial AML study showing that FTO demethylates m^6^A in mRNA^4^, while MOLM-13 cells were used in later studies^31,32^. Successful knockdown of FTO expression in both cell lines was confirmed by Western blot after generating stable cell lines by puromycin selection (≥5 days) (**Figure 1B**).

**Figure 1.**
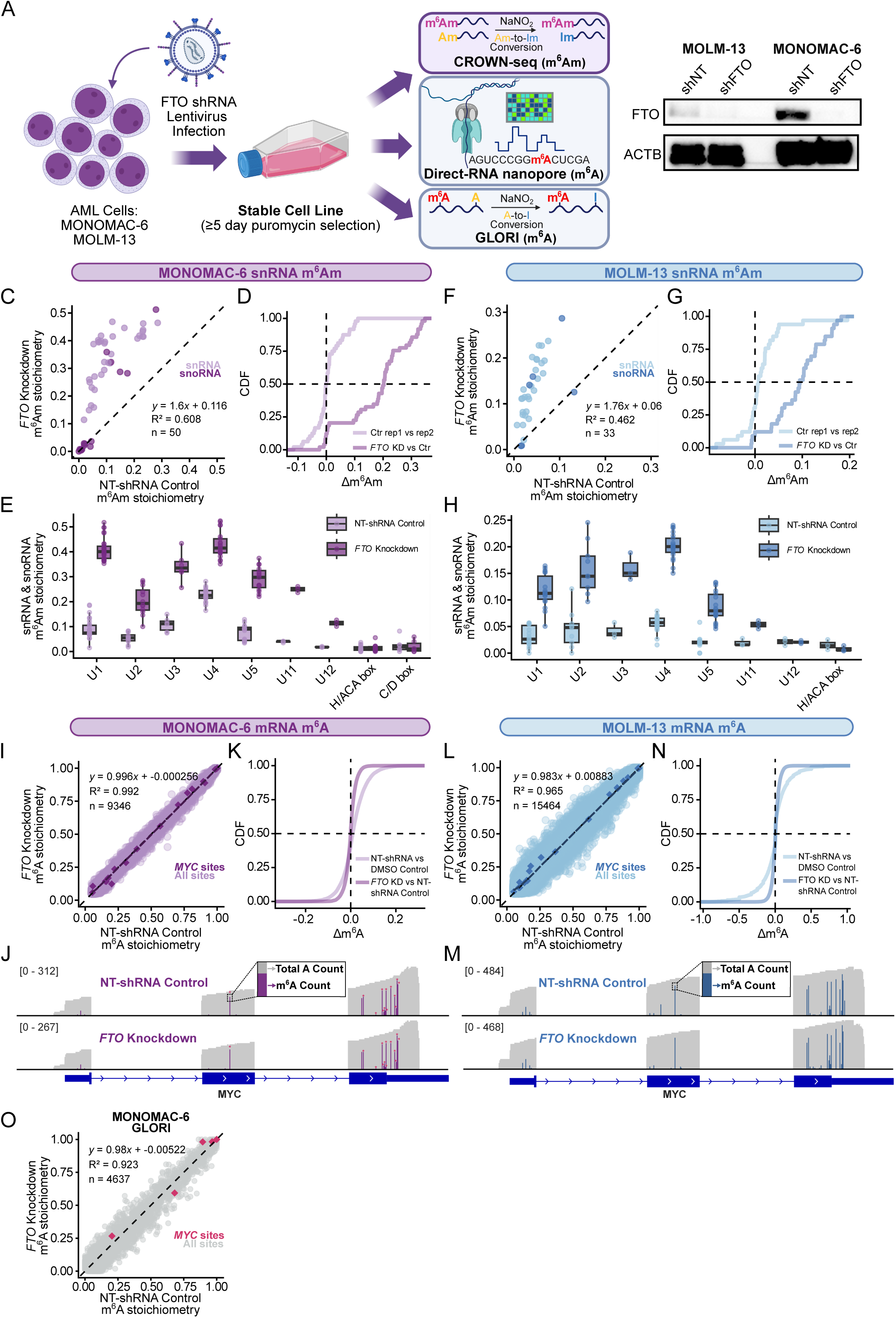
*FTO* knockdown in AML increases m^6^Am stoichiometry in snRNA but does not increase m^6^A stoichiometry in mRNA. **(A)** Schematic of lentivirus treatment and timing. Collected RNA was subjected to CROWN-seq for m^6^Am quantification, direct RNA nanopore-seq for m^6^A quantification, and GLORI for orthogonal m^6^A quantification. Created in BioRender. Jaffrey, S. (2026) https://BioRender.com/75xqx83. **(B)** Western blot assay confirming *FTO* knockdown by lentiviral constructs in MOLM-13 and MONOMAC-6 cells. ACTB (actin-ß) was used as a loading control. shNT, non-targeting shRNA control; shFTO, FTO-specific shRNA. **(C** and **F)** Scatter plots comparing snRNA and snoRNA transcription start site-specific m^6^Am stoichiometry in non-targeting control vs FTO-specific shRNA infected MONOMAC-6 and MOLM-13 cells, respectively. m^6^Am stoichiometry was quantified by CROWN-seq in triplicate and averaged for plotting. Only replicates with ≥50 reads per site were included. n, Pearson’s R^2^, and linear regression equation are denoted. **(D** and **G)** Cumulative distribution plots of m^6^Am differences (Δm^6^Am) in MONOMAC-6 and MOLM-13 cells, respectively. Included is a control comparison (i.e., comparison of NT-shRNA control replicate 1 vs replicate 2) to establish the normal replicate-to-replicate variability in this assay. Also shown is the experimental comparison (*FTO* shRNA knockdown vs NT-shRNA control, average of three replicates each), which demonstrates a substantial increase in stoichiometry of nearly all measured m^6^Am sites after FTO depletion in both cell lines. **(E** and **H)** Boxplots of m^6^Am stoichiometry in different snRNA and snoRNA transcript types in MONOMAC-6 and MOLM-13 cells, respectively. Three replicates each of NT-shRNA control and *FTO* knockdown show a robust increase in m^6^Am at every measured snRNA, as well as U3 snoRNA (only transcripts with ≥50 read coverage for each replicate are included). *FTO* knockdown resulted in an average increase of +0.235 across snRNAs in MONOMAC-6 cells, and +0.0932 in MOLM-13 cells. **(I** and **L)** Scatter plots comparing site-specific m^6^A stoichiometry between *FTO* knockdown and non-targeting shRNA control MONOMAC-6 and MOLM-13 cells, respectively (≥50 reads per site, m^6^A stoichiometry > 0.05). Each condition consists of three biological replicates merged for stoichiometry calculation. *MYC* sites are highlighted. n, Pearson’s R^2^, and linear regression equation are denoted. A LOESS curve is shown, which overlaps with the y=x identity line. **(J** and **M)** IGV (Integrated Genome Viewer) panels of *MYC* showing consistent per-site m^6^A stoichiometry in NT-shRNA control and *FTO* knockdown MONOMAC-6 and MOLM-13 cells, respectively (3 replicates merged for sufficient coverage). Comparable total read coverage is shown in grey. All DRACH sites were examined in both the control and *FTO* knockdown conditions. At each DRACH site, the fraction of A’s called as m^6^A is colored. For MONOMAC-6, eight sites that were also detected with GLORI in both conditions (≥25 reads per site) are denoted with a star above the corresponding site. **(K** and **N)** Cumulative distribution plots of m^6^A differences (Δm^6^A) in MONOMAC-6 and MOLM-13 cells, respectively. To establish the variation of m^6^A sites between replicates, we included a control comparison (NT-shRNA control vs DMSO control, three replicates merged), along with the experimental comparison (*FTO* knockdown vs NT-shRNA control, three replicates merged). No systematic shift is seen in these curves. The slightly larger variation in Δm^6^A in the control experiments reflects the lower read depth of these samples. **(O)** Scatter plot comparing site-specific m^6^A stoichiometry between *FTO* knockdown and NT-shRNA control MONOMAC-6 cells as measured by GLORI (≥50 reads per site, m^6^A stoichiometry > 0 in at least one condition). Each condition consists of three biological replicates merged for stoichiometry calculation. Five *MYC* sites are highlighted in pink. n, Pearson’s R^2^, and linear regression equation are denoted.

To confirm that FTO knockdown led to depletion of FTO activity, we analyzed m^6^Am levels in snRNA. As there is currently no nanopore basecaller for m^6^Am, we used CROWN-seq^11,33^, a single-nucleotide resolution method for stoichiometric mapping of m^6^Am. CROWN-seq uses the chemical reagent from GLORI, which converts 2’-*O*-methyladenosine (Am) to 2’-*O*-methylinosine (Im), but leaves *N*^6^,2’-*O*-dimethyladenosine (m^6^Am) intact^11,33^. By sequencing the transcription-start nucleotide and measuring the number of adenosines (m^6^Am) and inosines (Am), CROWN-seq provides highly accurate measurements of m^6^Am stoichiometry in snRNA^11,33^.

We compared m^6^Am stoichiometry at all annotated snRNA and snoRNA transcription-start site positions between three replicates of control and *FTO* knockdown conditions. All measured snRNA-associated m^6^Am sites, as well as U3 snoRNA m^6^Am sites, exhibited large increases in stoichiometry upon FTO depletion **(Figure 1C-H)**. In MONOMAC-6, the U1 snRNA gene family showed the largest increases in m^6^Am stoichiometry upon FTO depletion, with the methylated fraction increasing up to 0.347 (a 2.9-fold increase) compared to the control shRNA, indicating that U1 may be a mediator of FTO biology. In MOLM-13, the increases in stoichiometry were less pronounced, however, still robust, with the U4 gene family showing the largest increase of +0.157.

Overall, these data confirm that FTO is inhibited in the cells infected with the FTO shRNA-expressing lentivirus, based on both the western blot and the substantial increases in m^6^Am stoichiometry in snRNAs.

### FTO depletion does not affect the stoichiometry of m^6^A sites in AML mRNA

We next sought to identify the m^6^A sites in *MYC* mRNA that are demethylated by FTO in AML cells. We used the same RNA samples described above, comprising three biological replicates from the non-targeting control shRNA and three from the FTO shRNA-expressing MONOMAC-6 and MOLM-13 cells (**Figure 1A**). We subjected these samples to nanopore sequencing and m^6^A basecalling. We obtained 2.3-2.7 million primary mapped reads in both MONOMAC-6 and MOLM-13 cells, for both the FTO shRNA and non-targeting control shRNA lentivirus-infected cells (see **Table S1**).

To assess reproducibility across replicates, we generated pairwise scatter plots between individual biological replicates. These showed high correlation and similar transcriptome-wide m^6^A stoichiometry measurements between the biological replicates (**Figure S2A, B**).

We also examined the number of m^6^A sites that reached the 50 read threshold, both globally and for *MYC* (**Table S2**). The number of sites increased when merging the three biological replicates. Since the replicate-to-replicate reproducibility was high, we used the merged dataset to maximize the number of reads at each m^6^A site, thereby bringing more sites above the 50 read threshold and achieving the highest accuracy in calculating m^6^A stoichiometry at each site for each condition.

For our analysis, we measured m^6^A stoichiometry at every DRACH site in the transcriptome. We filtered out sites with stoichiometries less than 0.05. This resulted in a set of 9,346 global m^6^A sites and 30 *MYC-*specific m^6^A sites for MONOMAC-6 and a set of 15,464 global m^6^A sites and 39 *MYC-*specific m^6^A sites for MOLM-13, both of which include all m^6^A sites previously identified in *MYC* mRNA using GLORI^25^ (**Table S3**).

To identify the m^6^A sites in the *MYC* mRNA that increase in stoichiometry after FTO depletion, we prepared a scatter plot comprising the calculated stoichiometry of each detected m^6^A site throughout the transcriptome in the control shRNA and FTO-specific shRNA conditions (**Figure 1I, L**, and **S2C, D**). We first focused on the m^6^A stoichiometries for all DRACH sites in *MYC* (**Figure 1I, L**, highlighted). Surprisingly, we found that the stoichiometry was essentially identical in the FTO shRNA-expressing and control shRNA-expressing MONOMAC-6 cells at every m^6^A site in *MYC* mRNA. To better visualize the m^6^A sites in *MYC*, we compared IGV coverage tracks between conditions, with total read coverage shown in grey and m^6^A stoichiometry (represented as the fraction of modified reads) in purple or blue (**Figure 1J, M**). We did not identify FTO-associated changes at the surveyed *MYC* sites in our datasets.

Since we could not identify FTO-regulated m^6^A sites in *MYC*, we turned to other FTO-regulated transcripts reported to contribute to AML progression, including *CEBPA*^5^, *RARA*^4,6^, *ASB2*^4,6^, and *LILRB4*^34^ mRNA. Among these, only *CEBPA* achieved sufficient read coverage in our data (50 reads per site) to quantify m^6^A stoichiometry. Again, we saw no difference in m^6^A stoichiometry in FTO-depleted MONOMAC-6 and MOLM-13 cells relative to the control (**Figure S2E, F**). In the case of *RARA*, *ASB2*, and *LILRB4*, by relaxing the filter threshold to 10 reads, we could analyze several *LILRB4* and *RARA* m^6^A sites (**Figure S2G, H)**. Even in these noisier plots, none of these sites showed compelling evidence for increased stoichiometry in FTO-depleted cells.

Another possibility is that FTO is not transcript-specific, but induces global demethylation at all sites. However, we did not observe even a subtle global shift in m^6^A stoichiometries between control and FTO-depleted cells. The scatter plots showed tight correlations along the *y*=*x* identity line in both cell types **(Figure 1I, L).** Cumulative distribution plots of the difference in individual m^6^A site stoichiometries between *FTO* knockdown and control samples (Δm^6^A) confirm no apparent difference between knockdown and control in both MONOMAC-6 and MOLM-13 (**Figure 1K, N**). Furthermore, Bland-Altman analyses revealed mean differences and limits of agreement between FTO-depleted and control samples (**Figure S2I, J**) that were comparable to those observed between independent control samples (**Figure S1H, I**).

Overall, in contrast to m^6^Am which is readily demethylated by FTO in AML cells, our data suggest that FTO does not have a clear, robust, or readily detectable demethylase activity towards m^6^A on any transcript that is detected by nanopore sequencing.

### FTO depletion does not affect m^6^A stoichiometry in *MYC* using GLORI

Because of the unexpected absence of FTO-regulated m^6^A sites in the AML transcriptome, we decided to use GLORI to provide an orthogonal validation of the nanopore results. We generated GLORI libraries from the same RNA samples described above: three biological replicates each from non-targeting control shRNA and FTO shRNA-expressing MONOMAC-6 cells (**Figure 1A**). In total, we obtained 47.5 million primary mapped reads for the FTO-depleted MONOMAC-6 cells and 21.6 million reads for the control shRNA (see **Table S1**). GLORI measurements showed strong concordance with our nanopore dataset, with transcriptome-wide m^6^A stoichiometries highly correlated between the two methods, including at the detected *MYC* sites (**Figure S2K, L**).

In the GLORI dataset, five *MYC* m^6^A sites met the required coverage for quantification and are highlighted in pink (**Figure 1O**). As with the nanopore data, the *MYC* m^6^A sites showed no notable change in stoichiometry in FTO-deficient MONOMAC-6 cells. We also denote the locations of thirteen *MYC* sites with coverage ≥25 reads and m^6^A stoichiometry > 0.05 on the nanopore IGV plot (**Figure 1J**, pink stars). These sites include the major regulated site previously reported^5^ in exon 2, which we find is not increased in FTO-depleted cells (stoichiometry values of 1.0 in the control and 0.89 in the *FTO* knockdown) (see **Table S3**). We also detect numerous sites in the 3’ UTR, which were previously reported^5^ to have smaller increases upon FTO depletion, however our GLORI data does not reproduce this increase. Notably, these previously reported sites have very high stoichiometry in control shRNA conditions, making it impossible for their stoichiometry to increase markedly after FTO depletion.

We also asked if a global upward shift in stoichiometry could be detected in FTO-depleted cells, consistent with a non-specific demethylase function. However, as with nanopore sequencing, GLORI showed no systematic upward shift in m^6^A stoichiometry within our measurement precision in the *FTO* knockdown samples compared with the non-targeting controls (**Figure 1O**).

Overall, both quantitative mapping methods showed that FTO depletion does not lead to an appreciable increase in m^6^A stoichiometry in *MYC* or at other surveyed sites in the AML transcriptome.

### Pharmacologic inhibition of FTO using FB23-2 does not affect m^6^A stoichiometry in mRNA

As an alternative approach to test the role of FTO as an m^6^A demethylase, we inhibited FTO with FB23-2, a small-molecule inhibitor reported to upregulate global m^6^A levels and induce AML differentiation and apoptosis^6^. For these experiments, we treated MONOMAC-6 and MOLM-13 AML cells with 5 µM FB23-2 for 5 days before performing both direct RNA nanopore sequencing and CROWN-seq (**Figure 2A**). Three biological replicates for both drug and DMSO (vehicle) conditions were subjected to nanopore direct RNA sequencing and m^6^A basecalling. Again, the replicates showed high replicate-to-replicate similarity (**Figure S3A, B**). All replicates were merged for high-coverage single-nucleotide m^6^A stoichiometry calculation. In total, we obtained 1.7-3.8 million primary mapped reads in both MONOMAC-6 and MOLM-13 cells treated with FB23-2 or DMSO (see **Table S1**).

**Figure 2.**
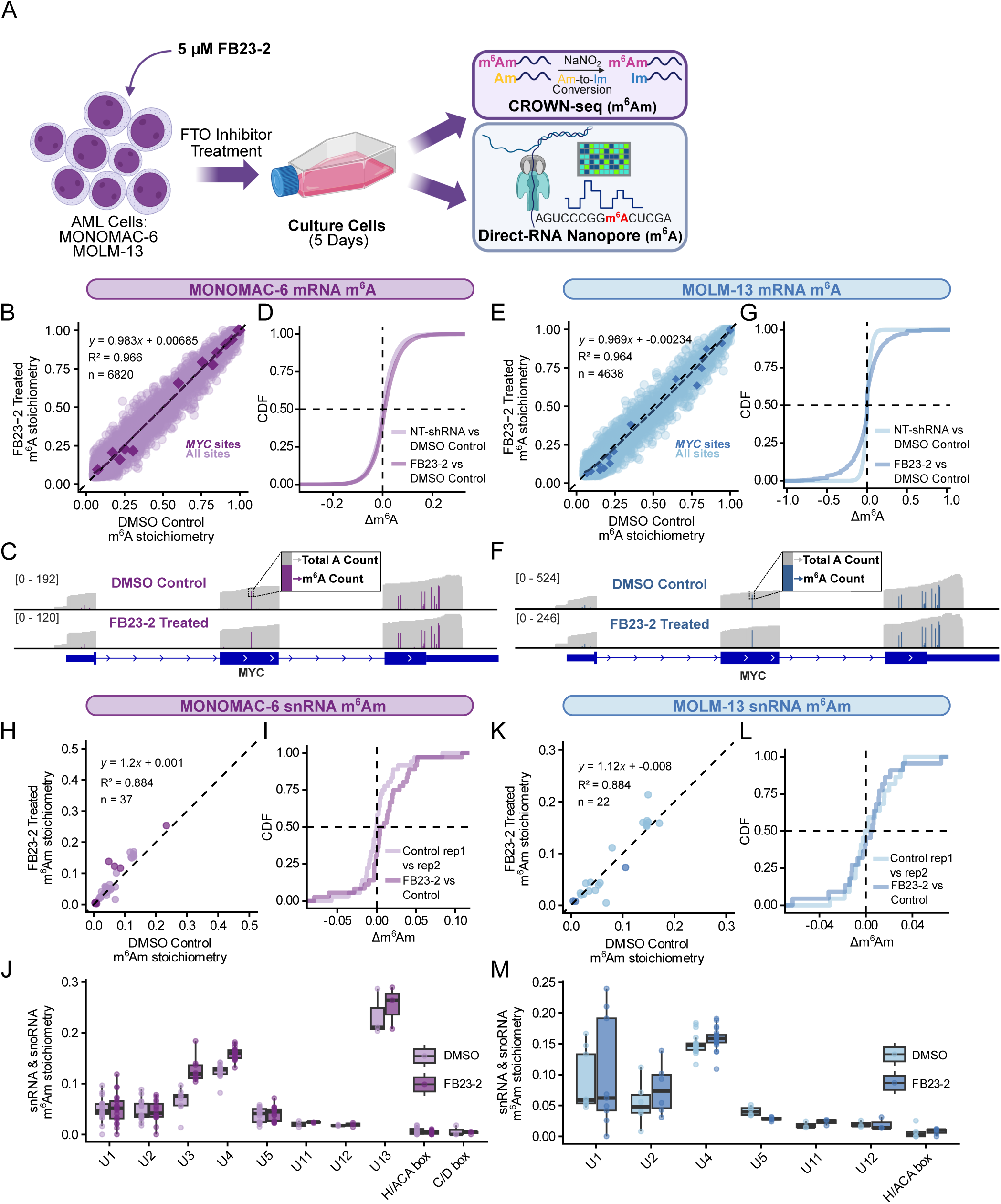
FTO inhibition via FB23-2 does not increase m^6^A stoichiometry in AML mRNA. **(A)** Schematic of FB23-2 drug treatment and timing. Collected RNA was subjected to direct RNA nanopore-seq for m^6^A quantification and CROWN-seq for m^6^Am quantification. Created in BioRender. Jaffrey, S. (2026) https://BioRender.com/0vo3cnzd. **(B** and **E)** Scatter plots comparing site-specific m^6^A stoichiometry between FB23-2-treated and DMSO-treated control MONOMAC-6 and MOLM-13 cells, respectively (≥50 reads per site, m^6^A stoichiometry > 0.05). Each condition consists of three biological replicates merged for stoichiometry calculation. *MYC* sites are highlighted. n, Pearson’s R^2^, linear regression equation, and LOESS curve are depicted. **(C** and **F)** IGV (Integrated Genome Viewer) panels of *MYC* showing consistent per-site m^6^A stoichiometry in DMSO-treated control and FB23-2-treated MONOMAC-6 and MOLM-13 cells, respectively (3 replicates merged for sufficient coverage). Total read coverage is shown in grey. At each DRACH site, the fraction of A’s called as m^6^A is colored. Fewer reads are seen in the FB23-2-treated samples, however, stoichiometry remains unchanged. **(D and G)** Cumulative distribution plots of m^6^A differences (Δm^6^A) in MONOMAC-6 and MOLM-13 cells, respectively. Control comparison (NT-shRNA control vs DMSO control, three replicates merged) and experimental comparison (FB23-2-treated vs DMSO-treated control, three replicates merged) are shown. **(H** and **K)** Scatter plots comparing snRNA and snoRNA transcription start site-specific m^6^Am stoichiometry in DMSO control vs FB23-2-treated MONOMAC-6 and MOLM-13 cells, respectively. m^6^Am stoichiometry was quantified by CROWN-seq in triplicate and averaged for plotting. Only replicates with ≥50 reads per site were included. n, Pearson’s R^2^, and linear regression equation are denoted. **(I** and **L)** Cumulative distribution plots of m^6^Am differences (Δm^6^Am) in MONOMAC-6 and MOLM-13 cells, respectively. Control comparison (DMSO control replicate 1 vs replicate 2) and experimental comparison (FB23-2 vs DMSO control, average of three replicates each) are shown. **(J** and **M)** Boxplots of m^6^Am stoichiometry in different snRNA and snoRNA transcript types in MONOMAC-6 and MOLM-13 cells, respectively. Three replicates each of DMSO control and FB23-2 treatment show small increases in certain classes (only transcripts with ≥50 read coverage for each replicate are included). In MONOMAC-6, small increases in m^6^Am stoichiometry were seen in U4 snRNA (+0.034 mean increase) and U3 snoRNA (+0.059 mean increase) **(J)**. In MOLM-13, small increases were seen in U2 snRNA (+0.023 mean increase) and U4 snRNA (+0.0089 mean increase) **(M)**. The m^6^Am stoichiometries of the remaining detected snRNA and snoRNA transcripts were not affected by 5µM FB23-2 treatment.

Scatter plots of transcriptome-wide m^6^A stoichiometries (FB23-2 vs DMSO) revealed high similarity at all measured sites between the two conditions in both cell types, including at *MYC* sites (**Figure 2B, E**, and **S3C, D**). Again, we included a *MYC* gene-level view with IGV tracks (**Figure 2C, F**). Even though the merged FB23-2 dataset showed a smaller total read coverage relative to the DMSO control, m^6^A stoichiometries remained consistent, with small differences in both directions within the bounds of normal variability. As such, we did not detect FTO-regulated *MYC* sites by FB23-2 inhibition. Additionally, we found that FB23-2 treatment did not produce a global increase in m^6^A stoichiometry across all measured sites, as evidenced by the scatter plots (**Figure 2B, E**) and cumulative distribution plots of m^6^A differences (**Figure 2D, G**). Bland-Altman analyses in both cell lines showed comparable mean differences and lower/upper limits of agreement (**FigureS3E, F**) relative to control comparisons **(Figure S1H, I**), suggesting no appreciable effect on stoichiometry beyond baseline variability.

We also assessed the effect of 5 µM FB23-2 on m^6^Am levels in snRNA by performing CROWN-seq on the same RNA input described above. The snRNA from the MONOMAC-6 samples exhibited subtle but detectable increases in m^6^Am (**Figure 2H, I**), primarily seen in the U3 snoRNA and U4 snRNA classes (**Figure 2J**). In MOLM-13 cells, however, there was no clear evidence of an increase in m^6^Am levels (**Figure 2K, L**), with only small increases seen in U2 and U4 snRNA classes (**Figure 2M**). The smaller effects of 5 µM FB23-2 on snRNA m^6^Am levels relative to our knockdown experiments may indicate that snRNA turnover does not occur during this time period, or that this concentration is insufficient to functionally inhibit FTO. Although increasing the concentration of FB23-2 would likely lead to more robust FTO inhibition and more robust increases in m^6^Am levels, as reported by Huang et al., higher concentrations resulted in markedly impaired cell proliferation^6^. During cell culture, we noted significant impairment of cell proliferation even at this mild concentration of 5 µM, which is discussed further below. We chose 5 µM for our studies since this was the concentration previously used to treat leukemia cell lines for measuring m^6^A changes using m6A-seq^6^.

Overall, using both the knockdown and pharmacologic conditions previously described to deplete FTO activity in AML cell lines, we find clear increases in snRNA m^6^Am levels by knockdown and subtler effects by drug inhibition. In contrast, we find no detectable increases in m^6^A stoichiometry in *MYC* or elsewhere in the transcriptome.

### Lack of selective FTO-regulation in non-AML cells

Although previous studies have highlighted the importance of FTO in mediating mRNA demethylation in AML, in part due to cytosolic localization of FTO in these cells^3^, we also examined other cell types. We recently characterized the effects of Fto depletion in mouse brain on behavior^35^. In these experiments, we prepared RNA from a basal forebrain region (nucleus accumbens shell, NAcSh) after stereotaxic injection of an AAV expressing Cre or GFP in 4-week-old *Fto*^fl/fl^ mice. After 5 weeks, Fto depletion was confirmed by *in situ* hybridization, and RNA was sequenced in 4 biological replicates for m^6^A quantification by nanopore direct RNA sequencing and m^6^Am quantification by CROWN-seq. In these experiments, we obtained ∼1 million reads cumulatively in both the Cre and GFP-expressing samples. We have reanalyzed the data here using a 50-read cutoff to quantify m^6^A stoichiometry after Fto depletion. Analysis of 7,846 shared m^6^A sites showed no evidence of elevated stoichiometry after Fto depletion at specific sites or globally across the transcriptome (**Figure 3A, B**). The scatter plot showed a tight correlation between conditions along the *y=x* identity line with no promising outliers, and the cumulative distribution of m^6^A differences showed no evidence of a rightward shift, suggesting that there was not even a subtle increase in global m^6^A stoichiometries upon Fto depletion. Bland-Altman analysis similarly showed no discernible change in m^6^A stoichiometry differences between conditions (**Figure S4A, B**).

**Figure 3.**
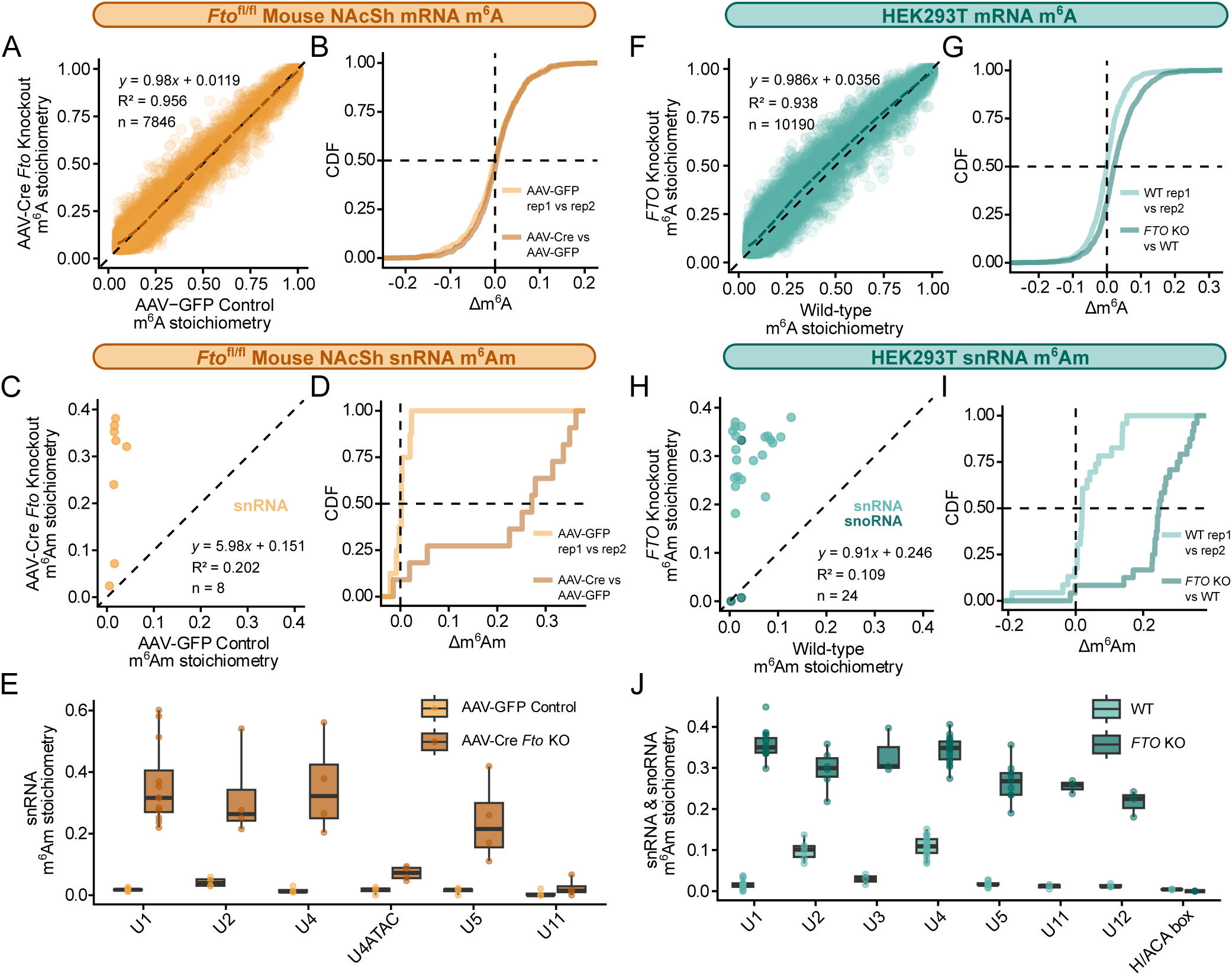
*FTO* knockout in non-AML cells increases m^6^Am stoichiometry in snRNA without affecting specific m^6^A site stoichiometries. **(A)** Scatter plot of site-specific m^6^A stoichiometries in the nucleus accumbens shell (NAcSh) of *Fto*^fl/fl^ mice treated with AAV-GFP control or AAV-Cre (≥50 reads per site, m^6^A stoichiometry > 0.05). Each condition consists of four biological replicates merged for stoichiometry calculation. n, Pearson’s R^2^, and linear regression equation are denoted. The LOESS curve shows no global upward shift in m^6^A stoichiometries after *Fto* conditional knockout. Data was previously deposited at GEO: <u>GSE334880</u>. **(B)** Cumulative distribution plot of m^6^A differences (Δm^6^A) in the NAcSh of *Fto*^fl/fl^ mice. Control comparison (AAV-GFP replicate 1 vs replicate 2) and experimental comparison (AAV-Cre vs AAV-GFP, four replicates merged) are shown. **(C)** Scatter plot comparing snRNA transcription start site-specific m^6^Am stoichiometry in AAV-GFP vs AAV-Cre *Fto*^fl/fl^ mouse NAcSh tissue. m^6^Am stoichiometry was quantified by CROWN-seq in quadruplicate and averaged for plotting. Only replicates with ≥50 reads per site were included. n, Pearson’s R^2^, and linear regression equation are denoted. Data was previously deposited at GEO: <u>GSE334881</u>. **(D)** Cumulative distribution plot of m^6^Am differences (Δm^6^Am) in the NAcSh of *Fto*^fl/fl^ mice. Control comparison (AAV-GFP replicate 1 vs replicate 2) and experimental comparison (AAV-Cre vs AAV-GFP, average of four replicates each) are shown. **(E)** Boxplots of m^6^Am stoichiometry in different snRNA transcript types in the NAcSh of *Fto*^fl/fl^ mice. Four replicates each of AAV-Cre *Fto* conditional knockout show a robust increase in m^6^Am in U1, U2, U4, and U5 snRNA (only transcripts with ≥50 read coverage for each replicate are included). *Fto* conditional knockout resulted in an average increase of +0.243 across snRNAs. Notably, m^6^Am levels rose from near zero to a max of 0.601 in U1 snRNA. **(F)** Scatter plot of site-specific m^6^A stoichiometries in WT vs *FTO* KO HEK293T cells (≥50 reads per site, m^6^A stoichiometry > 0.05). Each condition consists of three biological replicates merged for stoichiometry calculation. n, Pearson’s R^2^, and linear regression equation are denoted. The LOESS curve shows a slight global upward shift in m^6^A stoichiometries in the *FTO* KO condition. **(G)** Cumulative distribution plot of m^6^A differences (Δm^6^A) in HEK293T cells. Control comparison (wild-type replicate 1 vs replicate 2) and experimental comparison (*FTO* knockout vs wild-type, three replicates merged) are shown. The experimental comparison shows a subtle but clear rightward shift relative to the control comparison. **(H)** Scatter plot comparing snRNA and snoRNA transcription start site-specific m^6^Am stoichiometry in wild-type vs *FTO* knockout HEK293T cells. m^6^Am stoichiometry was quantified by CROWN-seq in triplicate and averaged for plotting. Only replicates with ≥50 reads per site were included. n, Pearson’s R^2^, and linear regression equation are denoted. **(I)** Cumulative distribution plot of m^6^Am differences (Δm^6^Am) in HEK293T cells. Control comparison (wild-type replicate 1 vs replicate 2) and experimental comparison (*FTO* knockout vs wild-type, average of three replicates each) are shown. **(J)** Boxplots of m^6^Am stoichiometry in different snRNA and snoRNA transcript types in HEK293T cells. Three replicates each of wild-type and *FTO* knockout show a robust increase in m^6^Am at every measured snRNA, as well as U3 snoRNA (only transcripts with ≥50 read coverage for each replicate are included). *FTO* knockout resulted in an average increase of +0.250 across snRNAs.

In contrast, we observed robust increases in m^6^Am in diverse snRNAs in the Cre-expressing tissue (**Figure 3C-E**). These effects were even larger than those seen in AML cell lines, with a mean increase of +0.242 and a max increase of +0.542. These data confirm that Fto activity was depleted in these tissues and further suggest that the primary effect of Fto is likely to be as an snRNA m^6^Am demethylase.

We were surprised that none of the cells or tissues tested thus far showed even subtle non-specific increases in m^6^A stoichiometry. Since FTO clearly can exhibit some m^6^A demethylase activity *in vitro*^2^, we expected a low level of m^6^A demethylase activity to be evident in the transcriptome. We suspected that this effect would require cells that exhibit higher FTO expression. We therefore considered HEK293T cells, which notably exhibit high levels of FTO protein relative to other cell types^11^, and exhibit highly efficient RNA processing activity, splicing, and translation^36^, which has contributed to its popularity for heterologous gene expression. Since snRNA and mRNA-regulatory pathways are highly expressed in HEK293T cells, we suspected these cells would be useful for detecting FTO activity on mRNA.

To test this, we analyzed wild-type and *FTO* knockout HEK293T cells^12^. Loss of FTO was reconfirmed by Western blot (**Figure S4C**), and replicates showed strong reproducibility in m^6^A measurements (**Figure S4D**). Here, when we examined the stoichiometries of m^6^A sites in WT and *FTO* knockout cells, we could observe a subtle but clear increase in m^6^A stoichiometry across all sites, as visualized by the LOESS curve (**Figure 3F, S4E**). Although a small global increase could be observed, no sites showed robust increases in stoichiometry, which would be expected if FTO targeted specific sites or mRNAs. Examining the cumulative distribution of m^6^A differences across all sites, we observed that the distribution was shifted in the positive direction by a fractional increase of +0.0211 (**Figure 3G**). Similarly, Bland-Altman analysis revealed that the mean difference in m^6^A stoichiometry rose from near-zero in the control comparison to +0.0314 in the *FTO* knockout vs WT comparison (**FigureS4F, G**). These data demonstrate that FTO can have a non-selective global effect on m^6^A stoichiometries, at least in HEK293T cells, although the magnitude of the effect is small.

In contrast to m^6^A, we again found marked increases in snRNA m^6^Am levels as measured by CROWN-seq, with an average increase in the methylated fraction of +0.250 (**Figure 3H-J**). Notably, relative to the AML cell lines, the m^6^Am levels in the FTO-depleted HEK293T cells were higher, and m^6^Am levels in the wild-type HEK293T cells were near-zero, suggesting elevated FTO activity in HEK293T cells, as expected.

### FTO is not required for AML cell line survival, and inhibitor FB23-2 retains cytostatic activity in FTO-deficient cells

During the course of our study, we unexpectedly observed that FTO depletion did not impair proliferation in our AML cell lines, in contrast to the previously published results^4–6^. The lack of an effect of FTO depletion on cell growth is inconsistent with FTO being a therapeutic target for AML. We therefore sought an independent assessment of whether FTO is required for AML cell line survival. For this, we queried the Cancer Dependency Map (DepMap), which integrates genome-wide CRISPR and RNAi loss-of-function screens to systematically identify genes required for cancer cell survival^37,38^. Established AML-essential genes like *MYB* and *CBFB*^39–42^ exhibited high dependency scores (>0.5), which defines a gene as essential for that particular cell type, across the 31 surveyed AML cell lines (**Figure S4H**). However, the dependency scores for *FTO* remained substantially below the 0.5 threshold (**Figure S4H**). Neither the CRISPR nor the RNAi dependency scores identified FTO as a required gene for AML cell line survival, suggesting that FTO is not an essential oncogenic dependency in leukemia cell culture.

Given that the DepMap does not identify FTO as an essential gene, it was confusing to us why the well-characterized FTO inhibitor FB23-2 would exert such potent anti-leukemic effects^6^. We hypothesized that the cytostatic activity of FB23-2 might arise from off-target mechanisms rather than direct FTO inhibition. To test this, we compared FB23-2-induced growth suppression in wild-type versus FTO-deficient cells. Strikingly, MTT viability assays revealed that FB23-2 suppresses proliferation in MONOMAC-6 cells lacking FTO just as effectively as it does in wild-type cells (**Figure S4I, J**). We observed a similar FTO-independent suppression of cell viability in non-leukemic HEK293T FTO-knockout cells (**Figure S4K, L**).

Together, these genetic and pharmacological data indicate that FTO is dispensable for AML cell line survival, and that the growth-inhibitory properties of FB23-2 are mediated by an alternative, FTO-independent cellular target. While FTO inhibitors may still possess therapeutic value through these alternative mechanisms, our findings caution against attributing suppression of cell growth by FB23-2 to functional proof of FTO’s oncogenic role or its epitranscriptomic regulatory activity.

## DISCUSSION

A core idea in epitranscriptomics is that FTO is an m^6^A eraser that removes methylation on specific transcripts, most notably *MYC*. The ability of FTO to demethylate *MYC* has popularized the idea of pharmacologic and genetic FTO inhibition as potential strategies to suppress AML^3–6,34^. The data supporting this model was generated with methods like MeRIP-seq and mass spectrometry that lack the ability to report the exact stoichiometry of m^6^A at single sites before and after FTO depletion. Using nanopore direct RNA sequencing and GLORI, two orthogonal methods capable of measuring absolute m^6^A stoichiometry at single sites, we found no detectable FTO-dependent increase in m^6^A at *MYC* or any other transcript in MONOMAC-6 and MOLM-13 AML cell lines, whether FTO was depleted by knockdown or pharmacologic inhibition. Similar results were seen outside of the AML context in *FTO* knockout brain and HEK293T cells. The only consistent, robust effect of FTO depletion was a marked increase in m^6^Am stoichiometry in snRNA and snoRNA, observed across every cell type and condition tested. Together, these data argue against a physiological role for FTO as a site-selective “m^6^A eraser” and instead point to m^6^Am in snRNAs as its primary substrate.

The discrepancy with earlier reports likely reflects methodology available at the time. Prior studies inferred FTO-regulated sites from increases in MeRIP-seq peak height^4–6^, which varies considerably between replicates even without any biological perturbation^16^, and, like other antibody-based enrichment methods such as ChIP-seq^43^, is not quantitative. A more accurate variant, m^6^A-seq2, exists^17^, and site-specific methods such as SCARLET^44^ and other quantitative mapping approaches^45–49^ are also available to quantify m^6^A stoichiometry at individual sites, but to our knowledge none of these have been applied to FTO depletion experiments. Thus, previous studies reported that FTO demethylated specific mRNAs without reporting the exact change in m^6^A stoichiometry after FTO depletion.

It is notable that our results differ from earlier reports of increased m^6^A/A by dot blot and mass spectrometry after FTO depletion^4–6,34^. Several studies have shown that bulk m^6^A/A measurements can be altered by global changes in transcript composition, not just site-specific methylation. In bladder cancer, for example, mass spectrometry showed that poly(A) RNA has reduced m^6^A compared to normal bladder^50^. However, subsequent GLORI analysis of individual m^6^A site stoichiometries did not show the expected site-specific drops in m^6^A stoichiometries. Instead, bladder cancer was found to exhibit increased expression of specific transcripts that have low levels of m^6^A transcripts, thus diluting the overall m^6^A levels in the transcriptome and causing the reduced m^6^A levels by mass spectrometry^50^. Similar transcriptomic alterations might explain m^6^A/A ratios after FTO depletion. Since FTO depletion shortens 3′UTRs by favoring proximal polyadenylation^51^, and because distal 3′UTR sequences are relatively m^6^A-poor, FTO depletion would be expected to lead to increased bulk m^6^A/A ratios. These confounding issues can affect bulk m^6^A measurements and underscore the need for site-specific, quantitative mapping for assigning FTO targets.

For quantitative mapping, we used direct RNA nanopore sequencing. The accuracy of nanopore sequencing is highly dependent on the settings used to interpret the data^20,22,23,26^, but with stringent modification probability filtering, the accuracy is quite high and matches the current state-of-the-art chemical conversion method, GLORI^25^. We favored nanopore sequencing over GLORI because it enables analysis of m^6^A in full-length transcripts. While we did not observe significant FTO-dependent differences in m^6^A stoichiometry, this approach remains valuable for future studies in which such changes might occur. In those scenarios, the full-length reads afforded by nanopore sequencing can distinguish between genuine FTO-mediated demethylation events from altered expression of different isoforms with distinct patterns of methylation.

A problem with the idea that FTO selectively demethylates specific m^6^A sites is somewhat difficult to understand. Since essentially all m^6^A sites share the same DRACH motif, it would be unclear how any single site would be selectively targeted. To achieve this, FTO would likely be selectively recruited by a second motif to a specific region, thus enabling it to demethylate proximal m^6^A sites. Indeed, artificial recruitment of FTO to specific sites in mRNA using a CRISPR-dCas13b-FTO fusion leads to efficient demethylation at proximal sites^52^. Although we did not detect evidence for such a mechanism, it remains possible that certain RNAs that were too lowly expressed to be detected have FTO-binding sites and could undergo selective demethylation. However, many of the specific mRNAs previously reported to be FTO targets were detectable and clearly did not show FTO-mediated demethylation, arguing against widespread selective demethylation of m^6^A by FTO.

If FTO does not selectively demethylate m^6^A, what is its function? We have found that m^6^Am is a substantially better FTO substrate than m^6^A^10–12^. In this study, snRNA and snoRNA m^6^Am stoichiometries rose dramatically after FTO depletion in every system tested, with mean baseline levels of 1.9-9.4% rising to 12.9-28.4% in FTO-deficient contexts. In HEK293T cells, which express relatively high FTO levels, we also detected a small, non-selective transcriptome-wide increase in m^6^A (∼2%), consistent with the weak demethylase activity previously reported for FTO *in vitro*^2^. However, it seems unlikely that the increase in fractional methylation of m^6^A by ∼2% would have a meaningful effect on transcript regulation. We therefore propose that the biological phenotypes of FTO depletion that were previously attributed to loss of m^6^A demethylation should be reconsidered as being mediated by elevated m^6^Am in snRNA, and thus altered snRNA functions.

A related puzzle is why FTO has been reported as required for AML survival if it does not regulate m^6^A at oncogenic transcripts. In our hands, FTO depletion did not impair proliferation of MOLM-13 or MONOMAC-6 cells, and DepMap CRISPR and RNAi screens across 31 AML cell lines likewise do not identify FTO as an essential gene^37,38^, in contrast to genes with established oncogenic roles in AML, such as *MYB* and *CBFB*^39–42^. We also found that the FTO inhibitor FB23-2 retained cytostatic activity in *FTO* knockdown MONOMAC-6 and *FTO* knockout HEK293T cells, indicating that its anti-proliferative effect is independent of FTO. While this study was in preparation, another group similarly reported that FB23-2 mediates cell proliferation suppression in an FTO-independent manner and showed that it targets DHODH, a key enzyme in uridine biosynthesis^53^. Together, these results suggest that FTO is dispensable for AML cell line survival in culture, and that the anti-leukemic activity previously attributed to FTO inhibition may reflect off-target effects.

Several limitations of this study should be noted. First, our stoichiometry measurements require a minimum read depth, so very lowly expressed transcripts were not captured. Second, nanopore basecalling accuracy is reduced near the 5′ end of reads^54^; improved 5′-ligation methods may address this^55^, though this limitation does not affect the *MYC* sites examined here, which are not located at the 5′ end of the transcript. Third, it is possible that m^6^A levels increase following FTO inhibition, but the affected mRNAs are degraded too rapidly to be detected. While this remains a possibility, the published studies did not report this effect. Instead, they found increased m^6^A levels after FTO inhibition, which should have been readily detectable by us using nanopore sequencing.

## ACKNOWLEDGEMENTS

We thank members of the Jaffrey lab for their comments and suggestions throughout the duration of this project. We thank the members of the Genomics core facility at Weill Cornell Medicine. We thank the Weill Cornell Medicine High-Performance Computing Core for providing computational resources (RRID:SCR_028969). We thank members of the Mason lab for valuable advice on nanopore sequencing and data processing, as well as for access to their PromethION sequencer. This work is supported by NIH grants R35 NS111631, S10 OD030335, RM1 HG011563, and R01 CA186702 (S.R.J.).

## AUTHOR CONTRIBUTIONS

L.S.N., C.G.T., and S.R.J. designed the experiments. L.S.N. and C.G.T. performed the experiments. J.F.L. analyzed CROWN-seq data. R.C and J.X. prepared *Fto*^fl/fl^ mouse brain samples. H.X.P. prepared and sequenced *Mettl3* KO samples. L.S.N. analyzed experimental data. L.S.N. and S.R.J. wrote the manuscript.

## SUPPLEMENTAL FIGURE LEGENDS

**Figure S1.**
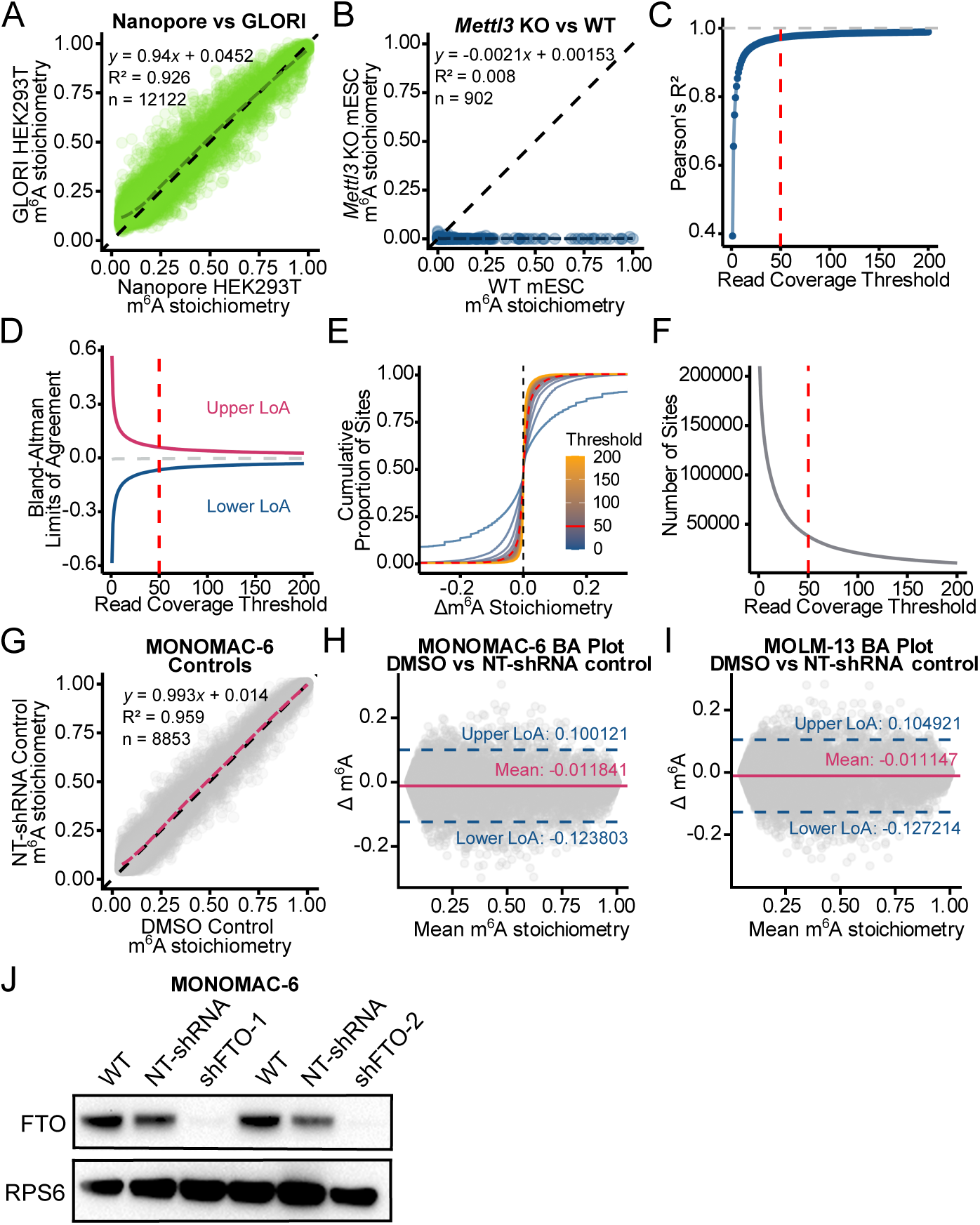
Justification of read depth threshold for m^6^A stoichiometry analysis. **Related to Figure 1. (A)** Confirmation of nanopore accuracy relative to GLORI. Scatter plot comparing site-specific m^6^A stoichiometry in HEK293T cells as measured by GLORI or nanopore direct RNA sequencing shows a robust correlation (≥50 reads per site, m^6^A stoichiometry > 0.05). GLORI data consists of 2 replicates merged from the public HEK293T dataset (GEO: <u>GSE210563</u>)^25^, and nanopore data consists of 3 replicates merged. Number of shared m^6^A sites (n), Pearson’s R^2^, and linear regression equation are shown. LOESS curve overlaid for visual trend estimation. **(B)** Confirmation of low false positive rate in nanopore basecalling. Scatter plot comparing nanopore site-specific m^6^A stoichiometry between wild-type and *Mettl3* KO mouse embryonic stem cells (≥50 reads per site, 1 replicate each). n, Pearson’s R^2^, linear regression equation, and LOESS curve are depicted. **(C)** Pearson’s correlation (R^2^) between m^6^A stoichiometry values obtained with two different control conditions: DMSO (3 replicates, merged) and control NT-shRNA (3 replicates, merged) in MONOMAC-6 cells. All samples were plotted against minimum read coverage threshold. Correlation strength begins to approach 1.0 at higher thresholds, indicating that stringent thresholding is required for accurate m^6^A stoichiometry calculation. At a threshold of 50, R^2^ = 0.973. **(D)** Upper and lower limits of agreement (LoA) between control samples plotted against minimum read coverage threshold. Limits of agreement were calculated by Bland-Altman analysis and demonstrate reduced variance in m^6^A stoichiometry measurements with higher thresholding. The mean per-site difference at each threshold is plotted as a grey, dashed line, demonstrating that increased thresholding reduces variance while the mean remains unaffected. At a threshold of 50, 95% of m^6^A differences between controls are expected to fall between –0.124 and 0.100. **(E)** Cumulative distribution plot of per-site m^6^A stoichiometry differences (Δm^6^A) between the above control samples for a series of minimum read coverage thresholds. All curves are centered near zero, indicating high concordance between controls. Increasing the read threshold produces steeper slopes and narrower distributions, reflecting reduced variance at higher read depths. The curve corresponding to a threshold of 50 is highlighted in red. **(F)** Number of shared, analyzable sites between control samples plotted against read coverage threshold. The number of sites decreased sharply as the threshold increased from zero, reflecting the exclusion of many low-coverage sites, and then began to plateau at higher thresholds as the remaining sites were supported by sufficient coverage. Paired with the previous analyses, we chose an optimal threshold of 50 for subsequent analyses to retain as many analyzable sites as possible. At a threshold of 50, the control samples had 38,018 shared sites. After further filtering for m^6^A stoichiometry > 0.05, the number of shared sites became 9,346. **(G)** Confirmation of nanopore reproducibility. Scatter plot comparing nanopore site-specific m^6^A stoichiometry between DMSO-treated and non-targeting shRNA-treated MONOMAC-6 cells (≥50 reads per site, m^6^A stoichiometry > 0.05). Each condition consists of three biological replicates merged for stoichiometry calculation. n, Pearson’s R^2^, linear regression equation, and LOESS curve are depicted. **(H** and **I)** Bland–Altman analysis comparing m^6^A stoichiometry measurements between DMSO and NT-shRNA control conditions in MONOMAC-6 and MOLM-13 cells, respectively. Each point represents the difference between paired measurements (DMSO control – NT-shRNA control) plotted against their mean. The solid horizontal line indicates the mean difference, while the dashed lines represent the limits of agreement (upper and lower LoA). This plot serves as a basis for the normal variance to be expected between two controls at a threshold of 50 reads minimum. **(J)** Western blot assay confirming *FTO* knockdown by two lentiviral constructs in MONOMAC-6 cells. RPS6 was used as a loading control. shNT, non-targeting shRNA control; shFTO-1, FTO-specific shRNA 1: TRCN0000246247; shFTO-2, FTO-specific shRNA 2: TRCN0000257473. shFTO-1 was chosen for downstream analysis and infection of MOLM-13 cells.

**Figure S2.**
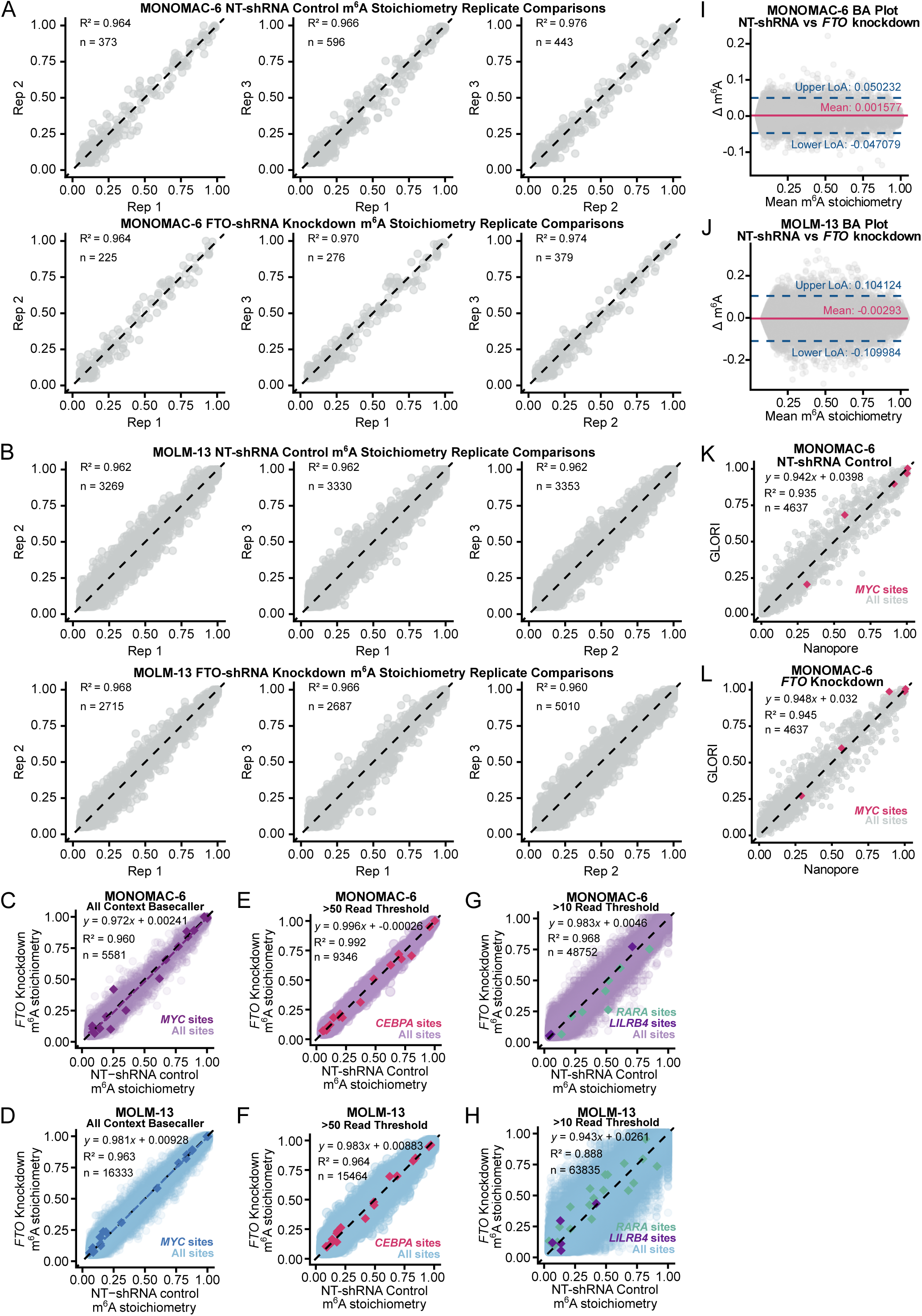
**Related to Figure 1. (A and B)** Pairwise scatter plots comparing per-site m^6^A stoichiometry calculations between three biological replicates in MONOMAC-6 and MOLM-13 cells, respectively (≥50 reads per site, m^6^A stoichiometry > 0.05). Non-targeting shRNA control replicates are compared in the top panel, and FTO-shRNA knockdown replicates are compared in the bottom panel. Pearson’s R^2^ and number of shared sites (n) are reported in each comparison. Note that more shared sites are retained in the MOLM-13 replicates, as each sample was sequenced at a higher read depth (see **Table S1**). **(C** and **D)** Scatter plots comparing site-specific m^6^A stoichiometry from the “All Context” basecaller between *FTO* knockdown and NT-shRNA control MONOMAC-6 cells (one replicate each) and MOLM-13 cells (three replicates merged), respectively (≥50 reads per site, m^6^A stoichiometry > 0.05). *MYC* sites are highlighted. n, Pearson’s R^2^, linear regression equation, and LOESS curve are depicted. To ensure that we are not missing any potentially FTO-regulated sites that lie outside of the DRACH consensus site context, we also basecalled our raw data using the “All Context” Dorado basecaller (inosine_m6A), in which every A nucleotide is assessed for modification probabilities. This result is very similar to the DRACH motif basecaller (see Figures 1H**, K**), with no sites exhibiting increased m^6^A stoichiometry outside the bounds of normal variability. **(E** and **F)** Scatter plots comparing site-specific m^6^A stoichiometry between *FTO* knockdown and NT-shRNA control MONOMAC-6 and MOLM-13 cells, respectively (≥50 reads per site, m^6^A stoichiometry > 0.05). *CEBPA* sites are highlighted in pink. n, Pearson’s R^2^, and linear regression equation are denoted. No *CEBPA* sites appear to be regulated by FTO. **(G** and **H)** Scatter plots with relaxed thresholding comparing site-specific m^6^A stoichiometry between *FTO* knockdown and NT-shRNA control MONOMAC-6 and MOLM-13 cells, respectively (≥10 reads per site, m^6^A stoichiometry > 0.05). *RARA* sites are highlighted in teal, and *LILRB4* sites are highlighted in purple. n, Pearson’s R^2^, and linear regression equation are denoted. No *RARA* or *LILRB4* sites appear to be regulated by FTO. **(I** and **J)** Bland–Altman analysis comparing m^6^A stoichiometry measurements between *FTO* knockdown and NT-shRNA control in MONOMAC-6 and MOLM-13 cells, respectively. Each point represents the difference between paired measurements (NT-shRNA control – FTO-shRNA) plotted against their mean. The solid horizontal line indicates the mean difference, while the dashed lines represent the limits of agreement. The mean differences and limits of agreement are comparable to the control comparisons (see **Figures S1G, H**). **(K** and **L)** Scatter plots comparing site-specific m^6^A stoichiometries in MONOMAC-6 cells measured by nanopore direct RNA sequencing versus GLORI sequencing (≥50 reads per site, m^6^A stoichiometry > 0 in at least one condition). *MYC* sites are highlighted in pink. Both the non-targeting shRNA control comparison **(K)** and *FTO* knockdown comparison **(L)** show good concordance between the two sequencing methods. n, Pearson’s R^2^, and linear regression equation are denoted.

**Figure S3.**
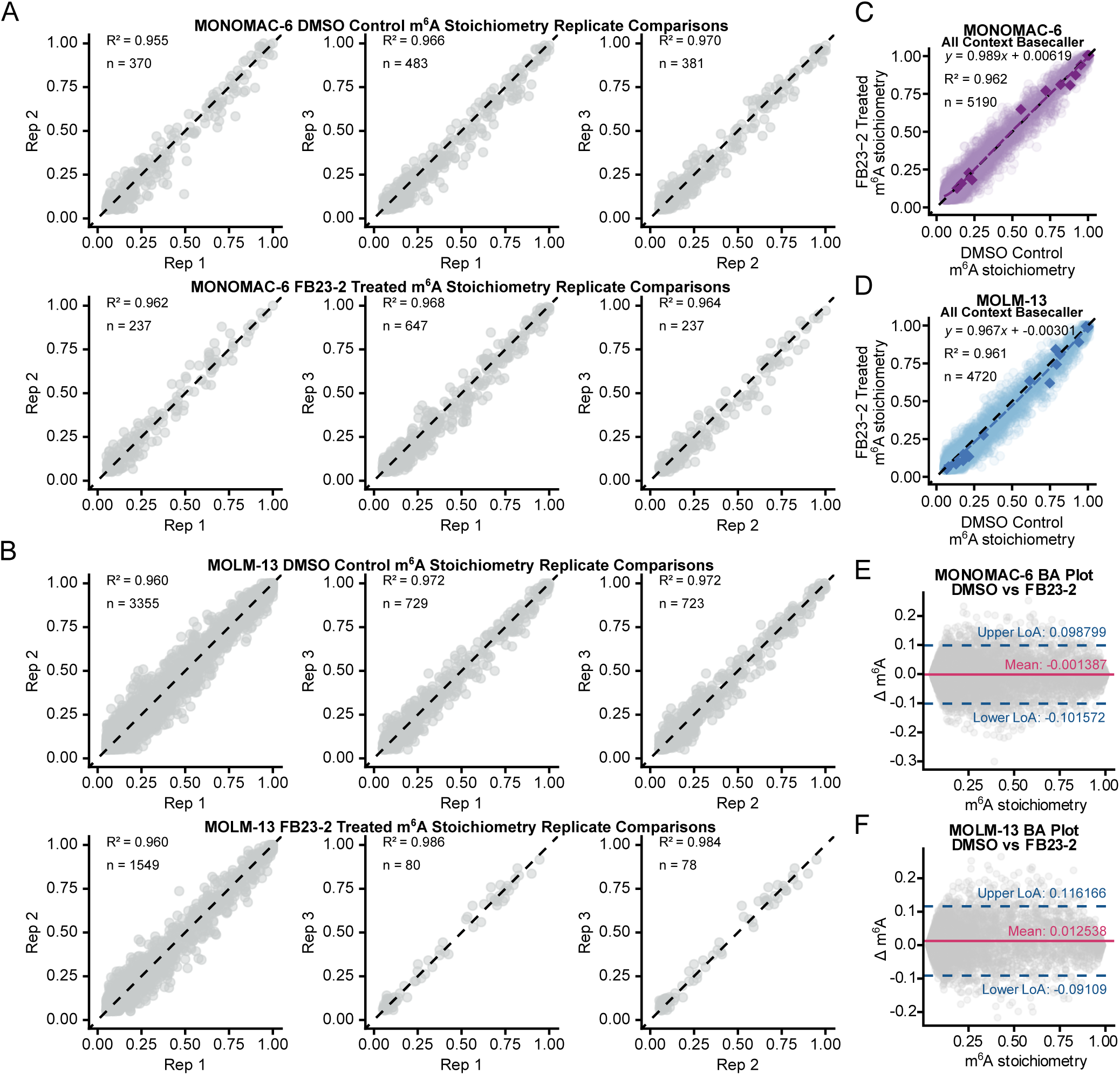
**Related to Figure 2. (A and B)** Pairwise scatter plots comparing per-site m^6^A stoichiometry calculations between three biological replicates in MONOMAC-6 and MOLM-13 cells, respectively (≥50 reads per site, m^6^A stoichiometry > 0.05). DMSO-treated control replicates are compared in the top panel, and FB23-2-treated replicates are compared in the bottom panel. Pearson’s R^2^ and number of shared sites (n) are reported in each comparison. **(C** and **D)** Scatter plots comparing site-specific m^6^A stoichiometry from the “All Context” basecaller between FB23-2-treated and DMSO-treated control MONOMAC-6 (1 replicate each) and MOLM-13 (three replicates merged) cells, respectively (≥50 reads per site, m^6^A stoichiometry > 0.05). *MYC* sites are highlighted. n, Pearson’s R^2^, linear regression equation, and LOESS curve are depicted. Similar to the knockdown experiments, we basecalled the raw data with the “All Context” basecaller model to assess non-DRACH m^6^A sites. The resulting scatter comparisons of FB23-2-treated and DMSO-treated control m^6^A sites show no sites exhibiting increased m^6^A stoichiometry. **(E** and **F)** Bland–Altman analysis comparing m^6^A stoichiometry measurements between FB23-2-treated and DMSO control MONOMAC-6 and MOLM-13 cells, respectively. Each point represents the difference between paired measurements (DMSO control – FB23-2-treated) plotted against their mean. The solid horizontal line indicates the mean difference, while the dashed lines represent the limits of agreement.

**Figure S4.**
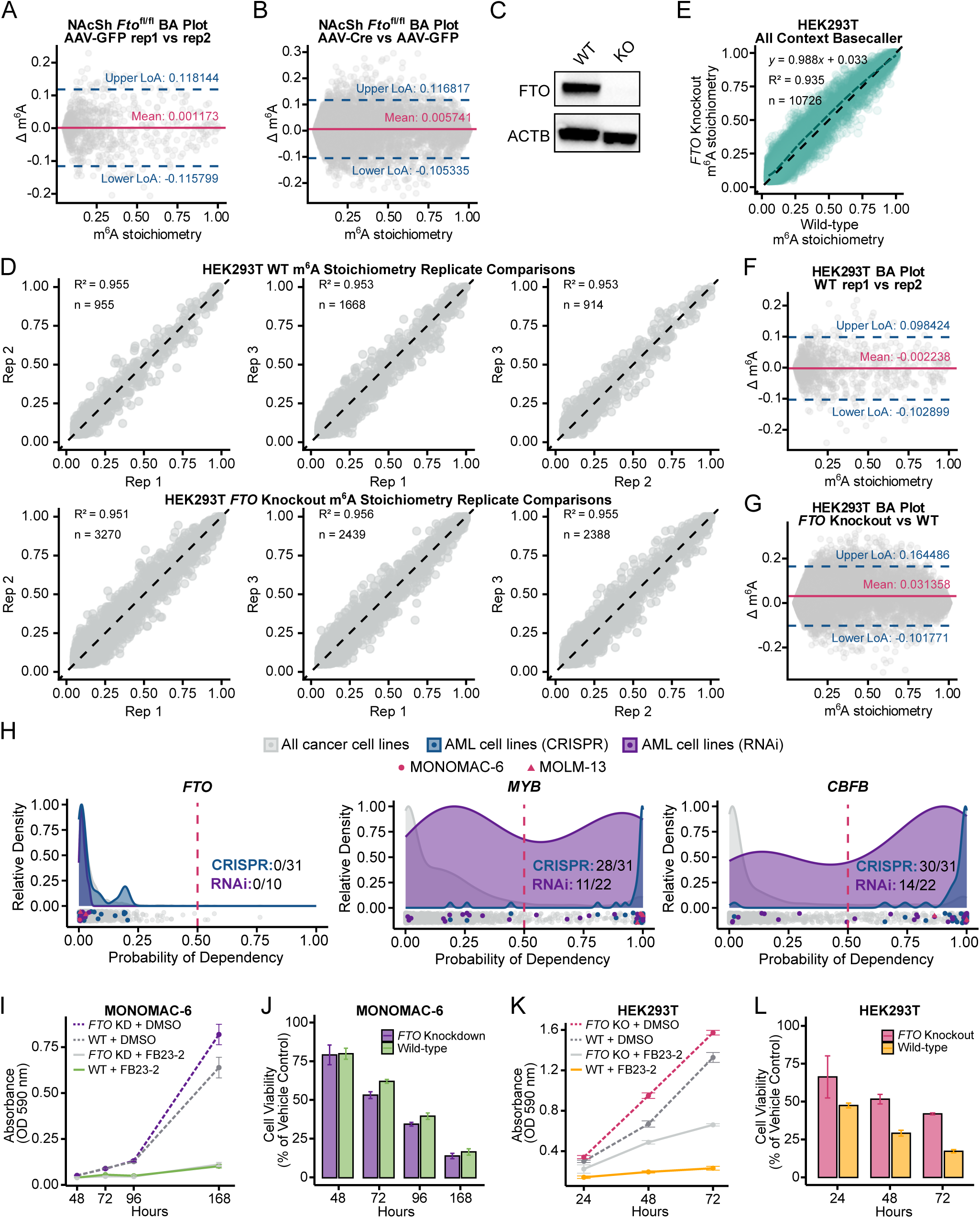
**Related to Figure 3. (A and B)** Bland–Altman analysis comparing m^6^A stoichiometry measurements in nucleus accumbens shell mouse tissue. The control analysis compares replicates of AAV-GFP control samples (replicate 1 vs replicate 2) **(A)**, and the experimental analysis compares AAV-Cre *Fto* knockout vs AAV-GFP control (four replicates merged) **(B)**. Each point represents the difference between paired measurements plotted against their mean. The solid horizontal line indicates the mean difference, while the dashed lines represent the limits of agreement. Note that the experimental comparison has more data points due to the merging of four replicates, nevertheless, the mean difference and limits of agreement do not substantially deviate from the control comparison. **(C)** Western blot assay reconfirming previously generated *FTO* knockout HEK293T cell line^12^. ACTB (actin-ß) was used as a loading control. WT, wild-type; KO, *FTO* knockout. **(D)** Pairwise scatter plots comparing per-site m^6^A stoichiometry calculations between three biological replicates in HEK293T cells (≥50 reads per site, m^6^A stoichiometry > 0.05). Wild-type replicates are compared in the top panel, and *FTO* knockout replicates are compared in the bottom panel. Pearson’s R^2^ and number of shared sites (n) are reported in each comparison. **(E)** Scatter plot comparing site-specific m^6^A stoichiometry from the “All Context” basecaller between *FTO* knockout and wild-type HEK293T cells (three replicates merged, ≥50 reads per site, m^6^A stoichiometry > 0.05). n, Pearson’s R^2^, linear regression equation, and LOESS curve are depicted. The “All Context” scatter plot closely matches the “DRACH” scatter plot (see Figure 4F). **(F** and **G)** Bland–Altman analysis comparing m^6^A stoichiometry measurements in HEK293T cells. The control analysis compares replicates of wild-type samples (replicate 1 vs replicate 2) **(F)**, and the experimental analysis compares *FTO* knockout vs wild-type (three replicates merged) **(G)**. Each point represents the difference between paired measurements plotted against their mean. The solid horizontal line indicates the mean difference, while the dashed lines represent the limits of agreement. Note that the experimental comparison has more data points due to the merging of three replicates. Here, we see that the experimental comparison diverges from the control comparison, with the mean difference and upper limit of agreement shifted slightly upwards. **(H)** The DepMap does not identify *FTO* (left) as an essential gene in most cell lines, but does identify known AML essential genes, *MYB* (middle) and *CBFB* (right). Density plots of Probability of Dependency scores from CRISPR (blue) and RNAi (purple) datasets show cell lines that are defined as dependent (Probability of Dependency score > 0.5). All cell lines are shown below each plot (grey), with AML cell lines highlighted in blue (CRISPR screen) or purple (RNAi screen), and MONOMAC-6 and MOLM-13 called out in pink. The number of AML cell lines defined as dependent for each gene is presented. Notably, no AML cell lines are considered to be dependent on *FTO*, and only 4 out of the 1183 cancer cells screened are found to be dependent. In contrast, most AML cell lines, including MONOMAC-6 and MOLM-13, are found above the 0.5 threshold and are defined as dependent on *MYB* and *CBFB,* as expected. **(I** and **J)** MTT assay of cell proliferation and cell viability time courses in wild-type and *FTO* knockdown MONOMAC-6 cells treated with DMSO or 5 µM FB23-2 (n = 6 replicates each). Cell proliferation (proportional to absorbance at OD=590 nm) was negatively affected by FB23-2 in both wild-type and *FTO* knockdown samples **(I)**. Cell viability, calculated as the percent absorbance in drug-treated over vehicle-treated control, showed a comparable reduction in both wild-type and FTO-depleted MONOMAC-6 cells after FB23-2 treatment **(J)**. Error bars represent mean ± standard error of the mean. **(K** and **L)** MTT assay of cell proliferation and cell viability time courses in wild-type and *FTO* knockout HEK293T cells treated with DMSO or 5 µM FB23-2 (n = 6 replicates each). Cell proliferation was negatively affected by FB23-2 in both wild-type and *FTO* knockout samples **(K)**. Cell viability showed a reduction in both wild-type and FTO-depleted HEK293T cells after drug treatment, with cells lacking FTO showing slightly higher viability **(L)**. Note that DMSO-treated samples reached 100% confluency at 72 hours. Error bars represent mean ± standard error of the mean.

## STAR Methods

### EXPERIMENTAL MODEL AND SUBJECT DETAILS

#### Cell lines

HEK293T (wild-type and *FTO* knockout cells, previously described by Mauer et al.^12^) were cultured in DMEM (Gibco, #11995065). MOLM-13 were grown in RPMI-1640 (ThermoFisher, 22400105). MONOMAC-6 were grown in RPMI-1640 supplemented with 2 mM L-Glutamine (ThermoFisher, 25030081), MEM NEAA (Life Technologies, 11140050), 1 mM sodium pyruvate (Life Technologies, 11360070), and 10 µg/ml human insulin (Life Technologies, 12585014). All media were supplemented with 10% fetal bovine serum (FBS) and 1X penicillin-streptomycin (Gibco, #15140148).

*Mettl3* KO and WT mESCs were previously described by Geula et al.^30^ and were a gift from S. Geula and J.H. Hanna (Weizmann Institute of Science). All mESCs were grown in gelatinized (0.1% gelatin in water, EmbryoMax ES-006-B) tissue culture plates in mESC media (KnockOut DMEM (Gibco #10829018), 15% heat-inactivated fetal bovine serum (FBS) (Gibco #26140079), 100 U/ml penicillin, 100 µg/ml streptomycin (Gibco #15140122), 1× GlutaMax (Gibco #35050061), 55 µM β-mercaptoethanol (Gibco #21985023), 1× MEM non-essential amino acids (Gibco #11140076), 1,000 U/ml LIF (Millipore ESG1107), 3 µM CHIR99201 (Sigma Aldrich SML1046), 1 µM PD0325901 (APExBIO A3013)).

Cells were incubated at 37L°C in a humidified atmosphere containing 5% CO_2_ and passaged as needed using TrypLE Express (ThermoFisher, 12604039).

#### Lentiviral constructs

pLKO lentiviral shRNA constructs and Non-Target control shRNA constructs were purchased from the Gene Editing & Screening (GES) Core Facility at MSKCC. Human FTO shRNA 1 cloneId: TRCN0000246247. Human FTO shRNA 2 cloneId: TRCN0000257473. Non-targeting control shRNA: TRC2 pLKO.5-puro Non-Target shRNA Control. Packaging plasmid: psPAX2(gag/pol/rev/tat). Envelope plasmid: pMD2.G (VSV-G).

### METHODS DETAILS

#### Lentiviral shRNA Knockdown

pLKO lentiviral shRNA constructs and Non-Target control shRNA constructs were purchased from the Gene Editing & Screening (GES) Core Facility at MSKCC (See **Table S4**). For lentiviral packaging, HEK293T cells were seeded in 100mm plates and allowed to reach 50-80% confluency. 2.5 µg of Lentiviral shRNA vectors were co-transfected with packaging plasmids (psPAX2 and pMD2.G) to each plate using FuGENE HD transfection reagent (Promega, E2311) according to the manufacturer’s instructions. Viral particles were harvested 24 hours, 48 hours, and 72 hours post-transfection, filtered through a 0.45 µm filter, pooled, and concentrated with Peg-it Virus Precipitation Solution (System Bioscience, #LV810A-1). 2×10^6^ MONOMAC-6 and MOLM-13 cells were suspended in a 12-well plate in 1 mL RPMI media containing 8 µg/ml polybrene (Santa Cruz Biotechnology, SC-134220). For each shRNA, a titration of virus (18µl:6µl:2µl) was used for infection. 48 hours post-infection, stable knockdown cells were selected with 1 µg/ml puromycin (Invivogen, ANT-PR-1) and cultured for an additional 5 days before collection in TRIzol LS or expansion of stable cell line. Given that each titration volume showed similar results, cells infected with 18 µl lentivirus were chosen for RNA and protein collection.

#### FB23-2 drug treatment

MONOMAC-6 and MOLM-13 cells were seeded in triplicate at 0.5×10^6^ cells/ml in 6-well plates. Cells were treated with 5 µM FB23-2 (MedChem Express LLC, #HY-127103) or 0.05% DMSO vehicle control and cultured for 5 days before collecting RNA in TRIzol LS.

#### RNA extraction and mRNA purification

Cellular total RNA in TRIzol LS (ThermoFisher #10296028) was extracted by Direct-zol RNA Miniprep kit (Zymo #R2070) or by Phenol Chloroform extraction. mRNA was purified by Dynabeads Oligo (dT)25 (Ambion #61002).

#### Cell Viability Assay

The effect of FB23-2 on cell viability was determined by 3-(4,5-dimethylthiazol-2-yl)-2,5-diphenyl tetrazolium bromide (MTT) assay. MONOMAC-6 (wild-type and *FTO* knockdown) cells were plated in 5 round-bottom 96-well plates with 200µl of 5×10^4^ cells/ml per well in 5 µM FB23-2 or 0.05% DMSO vehicle control and 6 replicates per condition. MTT assay was performed following Abcam’s instructions, with 5 timepoints: 24 h, 48 h, 72 h, 96 h, and 168 h. On each day, one plate was sacrificed by spinning down at 1,000 x g for 5 min at 4°C, followed by aspiration and resuspension in 50 µl serum-free RPMI media and 50 µl of 5 mg/ml MTT solution (Abcam, ab146345-500mg) in PBS. The plate was placed back in the incubator for 3 hours before adding 150 µl of MTT solvent (4 mM HCl, 0.1% NP40 in isopropanol) to dissolve formazan crystals by shaking on an orbital shaker for 15 min. Absorbance was read at OD = 590 nm, and DMSO vehicle-treated wells were considered as controls. Cell viability was calculated as:

Cell Viability (%) = (Mean absorbance of FB23-2 treated group/ Mean absorbance of control group) × 100

#### Western Blotting

Cells were lysed in lysis buffer [25 mM HEPES, pH 7.6, with 1.5 mM MgCl_2_, 420 mM NaCl, 0.5% NP-40, 0.2 mM EDTA, 25% (v/v) glycerol, 1 mM DTT, 1x protease inhibitor cocktail (ThermoFisher, 1861279)] with agitation for 30 min at 4°C. Supernatant was collected by spinning at 4°C at 16,000 × g for 20 min and quantified by Bradford assay (Bio-Rad, 5000006). 10 µg cell lysate was run on each lane of a NuPage 4-12% Bis-Tris pre-cast gel (ThermoFisher, NP0323BOX) and then transferred to a PVDF membrane (MilliporeSigma, IPVH00010). The membrane was blocked with 5% milk and blotted overnight at 4°C with 1:1000 anti-FTO rabbit monoclonal antibody (Abcam, ab124892), 1:2500 anti-RPS6 rabbit monoclonal antibody (Cell Signaling, 2217T), or 1:1000 anti-ACTB rabbit monoclonal antibody (Cell Signaling, 4970T) in 5% BSA/ TBST/ 0.1% NaN3 solution. 1:5000 HRP-conjugated anti-rabbit IgG antibody (Cytiva, NA934) was used as a secondary antibody in 5% milk for 1 h at room temperature. Immunoblots were developed using Amersham ECL Western Blotting Detection Reagent (Cytiva, RPN2209) on a Bio-Rad ChemiDoc Imaging system.

#### Nanopore direct RNA library preparation and sequencing

RNA sequencing libraries were prepared from total RNA or mRNA using the nanopore direct RNA Sequencing kit (ONT, SQK-RNA004) according to the manufacturer’s instructions, with some notable adjustments. Samples were multiplexed following the SeqTagger method from Pryszcz et al. 2025^23^ using the custom RT adapters for the b04_RNA004 and b96_RNA004 models (see **Table S4**). In brief, 500 ng total RNA or 100 ng polyA-selected RNA in 8 µl nuclease-free water were barcoded with 1µl custom RT adapters via ligation with 1.5 µl T4 DNA ligase (NEB, M0202L), 3 µl 5x NEBNext Quick Ligation reaction buffer (NEB, B6058S), 0.5 µl diluted RCS, and 1 µl SUPERase·In RNase Inhibitor (ThermoFisher #AM2696) for 15 min at room temperature. Samples were then reverse transcribed with 1 µl Induro Reverse Transcriptase (NEB, M0681L), 8 µl 5x Induro buffer, 2 µl 10 mM dNTP, and 14 µl nuclease-free water following the incubation protocol: 55°C for 50 min, 70°C for 10 min, and 12°C hold. RNA-cDNA products were cleaned up with 1.8X AMPure XP Beads (Beckman, A63881) and washed twice in 70% ethanol before eluting in water and pooling the 4 barcoded libraries together for a total of 20 µl. Next, the pooled library was ligated with 6 µl RNA ligation adapter (RLA), 3 µl T4 DNA ligase, 8 µl 5X NEBNext Quick Ligation buffer, and 3 µl nuclease-free water, incubating for 10 min at room temperature. The ligation product was cleaned up with 0.8X AMPure XP beads, washed twice with the provided wash buffer (WSB), and eluted in 33 µl RNA elution buffer (REB). Each library was quantified using 1 µl for Qubit 1x dsDNA HS assay (ThermoFisher, Q33231), with a recovery aim greater than 30ng total. Samples were loaded onto primed MinION (ONT, FLO-MIN004RA) or PromethION flow cells (ONT, FLO-PRO004RA) and sequenced on a MinION or PromethION for 2-24 hours, acquiring 150 Mb to 11.1 Gb (see **Table S1** for details).

#### Flow Cell Nuclease Flushing

For Flow Cells which were washed and reused, a modified Flow Cell wash mix^23^ was prepared with 20 µl TURBO DNase (2U/µl) (Thermo Fisher, AM2238) and 380 µl DIL (wash diluent, provided with ONT EXP-WSH004). Wash mix was loaded onto Flow Cell and incubated for 20 min before storing or loading another library.

#### Nanopore Data Analysis

The raw POD5 sequencing data files were basecalled using Dorado v1.0.0 with the rna004_130bps_sup@v5.2.0 m6A_DRACH modification model (for supplementary analysis, the rna004_130bps_sup@v5.1.0 inosine_m6A model was used, with m^6^A calls extracted). The raw POD5 files were also used to generate a demultiplexed pod5.demux.tsv.gz file following the SeqTagger protocol with model b04_RNA004 or b96_RNA004^23^. Basecalled BAM files were mapped to the human genome hg38 (or mouse genome mm10) using the Dorado aligner with Minimap2^56^ options “-ax splice -k 14”. Aligned reads were demultiplexed with the SeqTagger bam_split_by_barcode.py script using the previously generated pod5.demux.tsv.gz file^23^. The demultiplexed reads were sorted and indexed with SAMtools^57^ v1.2.20 and visualized using IGV^58^ v2.18.2 (a base modification likelihood threshold of 0.99 was used for visualizing m^6^A calls). For increased read coverage, replicates were merged with SAMtools merge before being sorted and indexed. Modification bed tables (mod.bed) were compiled using Modkit v0.2.8 with a pileup filter threshold of 0.99 for all bases, ensuring that only high-confidence calls are used for analysis. Mod.bed files were loaded into RStudio v2024.12.1+563 for subsequent analysis and figure drawing by a custom script.

#### snRNA CROWN-seq library preparation and sequencing

To study snRNA m^6^Am stoichiometries, we used a modified protocol of CROWN-seq^11^, in which 2’-*O*-methyladenosine (Am) is chemically converted into 2’*O*-methylinosine (Im) and *N*^6^,2’*O*-dimethyladenosine (m^6^Am) remains unconverted, allowing for m^6^Am quantification. Short RNAs (17-200 nt) were isolated from total RNA using RNA Clean & Concentrator-5 (Zymo Research, R1016), following the Purification of Small and Large RNAs into Separate Fractions protocol. 100 ng of the small RNA fraction, in 8 µl nuclease-free water, was used as input. Following the conditions from the GLORI2.0 protocol^59^, samples were converted in deamination buffer (13μl glyoxal, 10μl DMSO, 10μl 5LM NaNO_2_, 5μl H_3_BO_3_, 4μl 500LmM MES (pH 6.0)) for 10 min at 50°C. Next, samples were ethanol precipitated, and the glyoxal adduct was removed with deprotection buffer (500 mM TEAA pH=8.6, 47.5% deionized formamide, 2mM EDTA) for 10 min at 95°C. RNA was collected again by ethanol precipitation, denatured for 2 min at 65°C, and spiked with 10 ng of Drosophila melanogaster embryo Poly A+ RNA (Takara Bio, #636224) for QC. To eliminate contamination of RNA species with a 5’-triphosphate or 5’-monophosphate, a dephosphorylation reaction was set up with 5 µl 10X CutSmart buffer, 5 µl Quick CIP (5 U/µl) (NEB, M0525L), and 1 µl SUPERase·In RNase inhibitor (ThermoFisher #AM2696) for 2 hours at 37 °C. The reaction was cleaned up using a Zymo RCC-5 column, and the RNA was eluted in 15.5 µl water. Next, RNA was denatured for 5 min at 65°C before removing the m^7^G-cap with 2 µl mRNA Decapping Enzyme (100U/µl) (NEB, M0608S), 2 µl 10X Decapping Enzyme Reaction Buffer, and 0.5 µl SUPERase·In RNase inhibitor for 1 hour at 37°C. The decapped 5’-monophosphate RNA was purified with ethanol-AMPure XP (RNA:beads:ethanol=1:2:3), washed twice in 80% ethanol, and eluted in 11.5 µl water. 5’ ligation was carried out with 20 pmol of 5’ adapter (see **Table S4**), 2 µl T4 RNA ligase 1 (ssRNA Ligase, 30U/µl) (NEB, M0437M), 4 µl 10X T4 RNA Ligase Buffer, 4 µl 10mM ATP, 0.5 µl SUPERase·In RNase inhibitor, and 16 µl 50% PEG8000 for 2 hours at 25 °C. 5’ ligation product was again purified with ethanol-AMPure XP and eluted in 16 µl water. End repair was carried out with 1 µl T4 PNK (NEB, M0201L), 2 µl 10X T4 PNK Reaction Buffer, and 1 µl SUPERase·In RNase inhibitor for 30 min at 37 °C. End-repaired RNA product was purified with ethanol-AMPure XP and eluted in 12 µl water. 3’ adapter incorporation and cDNA synthesis were carried out via a template switching reaction with an R2R RNA/R2R DNA heteroduplex adapter based on TGIRT-seq^60–62^ (since A is assumed to be converted from GLORI, the R2R DNA mix consists of A, G, or C overhangs in a 1:1:2 ratio, see **Table S4**). The template switching reaction was first pre-incubated at room temperature for 30 min with 2 µl 10X annealed R2 RNA/R2R DNA heteroduplex mix (100 nM final conc.), 1 µl Induro RT (200U/µl) (NEB, M0681L), 4 µl 5X Induro RT Buffer, and 1 µl SUPERase·In RNase inhibitor; next, 1 µl 10mM dNTP was added before incubating for 5 min at 25 °C, 15 min at 55 °C, 1 min at 95 °C, and holding at 4 °C. 5 µl of the cDNA product was carried forward for indexing PCR, mixed with 15 µl water, 25 µl Phusion High-Fidelity PCR Master Mix with HF Buffer (NEB, M0531L), 2.5 µl i5 indexing primer, and 2.5 µl i7 indexing primer (NEB, E7780S). Cycling conditions were 98 °C for 30 sec, then 14 cycles of 98 °C 15 seconds, 65 °C 30 seconds, and 72 °C 15 seconds, with a final 5 min 72 °C extension. Two rounds of 0.9X AMPureXP bead purifications were performed to remove primers. 10-20 ng indexed library was obtained for each library. The libraries were mixed and sequenced by NovaSeqX.

#### snRNA CROWN-seq data processing

The read pairs were first quality trimmed by Cutadapt v3.4^63^: -m 32 -q 20 -e 0.25 -a NAGATCGGAAGAGCACACGTC -A TAANNNNNNNNNNNGATCGGAAGAGCGTCGTG. The subsequent alignment and statistics were identical to the CROWN-seq mapping pipeline as previously described^11^. In brief, the UMIs (the 11 nt at the beginning of read1) were extracted by UMI-tools v1.1.1^64^. Next, *in silico* converted read pairs (read1 A-to-G, read2 T-to-C) were aligned by HISAT2^65^ against the respective converted reference genome and transcriptome.

Only read1 reads without 5’ end softclips were used for 5’ end analysis. Pileup was performed to obtain the read coverages of every 5’ end in the transcriptome. Non-conversion rates of the transcription start nucleotides were calculated by A counts over A and G counts. Only canonical snRNA TSSs based on annotations from Gencode v45 were considered for our analysis.

#### GLORI library preparation and sequencing

To study mRNA m^6^A stoichiometries, we used a modified protocol of GLORI 2.0^59^. mRNA was first purified from 1 µg of total RNA by one round of Poly(A) selection by Dynabeads Oligo (dT)25 (Ambion #61002). mRNA was fragmented into 150-200nt fragments by incubation at 94°C for 3 min with 2µl 10X RNA Fragmentation buffer (NEB E6150S); the reaction was terminated with 2µl 10X stop buffer and cleaned-up by ethanol precipitation. Prior to conversion, samples were denatured at 65°C for 5 min and then stored on ice to prevent secondary structures from impeding conversion. Following the conditions from the GLORI2.0 protocol^59^, samples were converted in deamination buffer (13μl glyoxal, 10μl DMSO, 10μl 5LM NaNO_2_, 5μl H_3_BO_3_, 4μl 500LmM MES (pH 6.0)) for 10 min at 50°C. Next, samples were ethanol precipitated, and the glyoxal adduct was removed with deprotection buffer (500 mM TEAA pH=8.6, 47.5% deionized formamide, 2mM EDTA) for 10 min at 95°C. RNA was collected again by ethanol precipitation, and end repair was carried out with 1 µl T4 PNK (NEB, M0201L), 2 µl 10X T4 PNK Reaction Buffer, and 1 µl SUPERase·In RNase inhibitor (ThermoFisher #AM2696) for 30 min at 37 °C. The reaction was cleaned up using an RNA Clean & Concentrator-5 column (Zymo Research, R1016), and the RNA was eluted in 10µl water. 5’ ligation was carried out with 20 pmol of 5’ adapter (see **Table S4**), 2 µl T4 RNA ligase 1 (ssRNA Ligase, 30U/µl) (NEB, M0437M), 4 µl 10X T4 RNA Ligase Buffer, 4 µl 10mM ATP, 0.5 µl SUPERase·In RNase inhibitor, and 16 µl 50% PEG8000 for 2 hours at 25 °C. 5’ ligation product was purified with ethanol-AMPure XP (RNA:beads:ethanol=1:2:3), washed twice in 80% ethanol, and eluted in 10 µl water. 3’ adapter incorporation and cDNA synthesis were carried out via a template switching reaction with an R2R RNA/R2R DNA heteroduplex adapter based on TGIRT-seq^60–62^ (see **Table S4**). The template switching reaction was first pre-incubated at room temperature for 30 min with 2 µl 10X annealed R2 RNA/R2R DNA heteroduplex mix (100 nM final conc.), 1 µl Induro RT (200U/µl) (NEB, M0681L), 4 µl 5X Induro RT Buffer, and 1 µl SUPERase·In RNase inhibitor; next, 1 µl 10mM dNTP was added before incubating for 5 min at 25 °C, 15 min at 55 °C, 1 min at 95 °C, and holding at 4 °C. 5 µl of the cDNA product was carried forward for indexing PCR, mixed with 15 µl water, 25 µl Phusion High-Fidelity PCR Master Mix with HF Buffer (NEB, M0531L), 2.5 µl i5 indexing primer, and 2.5 µl i7 indexing primer (NEB, E7780S). Cycling conditions were 98 °C for 30 sec, then 15 cycles of 98 °C 15 seconds, 65 °C 30 seconds, and 72 °C 15 seconds, with a final 5 min 72 °C extension. Two rounds of 0.9X AMPureXP bead purifications were performed to remove primers. 2-20 ng indexed library was obtained for each library. The libraries were pooled, purified one more time by 1.0X SPRIselect beads (Beckman Coulter, B23318), and sequenced by NovaSeqX.

#### GLORI data processing

GLORI data processing was performed as previously described by us^11^. In brief, reads were first quality trimmed by Cutadapt v3.4^63^: -m 32 -j 4 -q 20 -e 0.25 -a NAGATCGGAAGAGCACACGTC -A TAANNNNNNNNNNNGATCGGAAGAGCGTCGTG. After pre-processing, the reads were A-to-G converted and aligned to converted reference genome and transcriptome by HISAT2 v2.1.0^65^: -k 5 –fr –rna-strandness FR –no-temp-splicesite –no-mixed. Only unique alignments were used. After alignment, a customized script was used to pileup every single base to obtain the A, C, G, and U counts. Non-conversion rate was calculated by A counts over A and G counts. Filters were applied to obtain high-quality non-converted A (m^6^A) signals. Details of this pipeline can be found at https://github.com/jhfoxliu/GLORI_pipeline.

#### DepMap Analysis

CRISPR gene dependency probability scores were obtained from the DepMap Public 25Q2 dataset^66^, and RNAi gene dependency probability scores were obtained from the 2019 DEMETER 2 Combined RNAi dataset^67^. We extracted the gene dependency probability scores of 1,183 cell lines for the genes *FTO, MYB,* and *CBFB* and plotted them to identify dependent cell lines. Gene dependency probability scores greater than 0.5 indicate that a cell line is dependent on the gene.

#### Genomic assembly and annotations

The following annotations and genomic sequences were used in this study: human, Gencode v45 (GRCh38); mouse, Gencode M10 (GRCm38.p4)

### QUANTIFICATION AND STATISTICAL ANALYSIS

Statistical analysis for CROWN-seq and GLORI were performed with Python (version 3.8.7). All other statistical analyses and visualizations were performed in RStudio Server (Posit Software, version 2024.12.1) running R version 4.3.3 (2024-02-29)^68^. Correlations between m^6^A stoichiometry values were assessed using Pearson’s product–moment correlation (cor.test function in R), with the coefficient of determination (R²) reported. Linear regression models were fitted with the lm function to calculate regression slope and intercept values. Bland–Altman analysis was performed to assess agreement between samples, with the mean difference and 95% limits of agreement (mean difference ± 1.96 × standard deviation of differences) reported. Exact R², linear regression equations, and Bland-Altman values are provided in each plot.

Versions of key Python packages: numpy (1.23.5); pandas (1.5.2); scipy (1.9.3); matplotlib (3.6.2); seaborn (0.12.1); matplotlib-venn (0.11.9). Versions of key R packages: ggplot2 (3.5.1); tidyr (1.3.1); dplyr (1.1.4); scales (1.3.0); patchwork (1.3.1)

## DATA AND CODE AVAILABILITY

All sequencing data generated from this study can be accessed from NCBI Gene Expression Omnibus (GEO): <u>GSE311653</u> for nanopore direct RNA sequencing data; <u>GSE311676</u> for CROWN-seq data; and <u>GSE311735</u> for GLORI data. The mouse nucleus accumbens shell sequencing data^35^ reanalyzed in this study were previously deposited at GEO under accessions <u>GSE334880</u> and <u>GSE334881</u>. The public GLORI HEK293T dataset was downloaded from GEO under accession code <u>GSE210563</u>. The public 25Q2 DepMap dataset is available from the Broad Institute’s DepMap Consortium and is accessible at https://depmap.org/portal/download/. Raw direct RNA nanopore data (POD5 files) are available upon request. Code for statistical analyses and illustrations is available upon request.

## MATERIALS AVAILABILITY

Reagents and materials in this study are available upon request.

## SUPPLEMENTAL INFORMATION

Table S1. Nanopore and GLORI sequencing information. Related to Figures 1, 2, and 3.

Table S2. Total DRACH and m^6^A sites per sample. Related to Figures 1, 2, and 3.

Table S3. *MYC* m^6^A sites in nanopore samples compared to GLORI. Related to Figures 1 and 2.

Table S4. shRNA constructs and sequencing adapters. Related to Figures 1, 2, and 3.

